# Defining the EM-signature of successful cell-transfection

**DOI:** 10.1101/2024.03.07.583927

**Authors:** Joshua G. Pemberton, Tatyana Tenkova, Philip Felgner, Josh Zimmerberg, Tamas Balla, John Heuser

## Abstract

In this report, we describe the architecture of Lipofectamine 2000 and 3000 transfection- reagents, as they appear inside of transfected cells, using classical transmission electron microscopy (EM). We also demonstrate that they provoke consistent structural changes after they have entered cells, changes that not only provide new insights into the mechanism of action of these particular transfection-reagents, but also provide a convenient and robust method for identifying by EM which cells in any culture have been successfully transfected. This also provides clues to the mechanism(s) of their toxic effects, when they are applied in excess. We demonstrate that after being bulk-endocytosed by cells, the cationic spheroids of Lipofectamine remain intact throughout the entire time of culturing, but escape from their endosomes and penetrate directly into the cytoplasm of the cell. In so doing, they provoke a stereotypical recruitment and rearrangement of endoplasmic reticulum (ER), and they ultimately end up escaping into the cytoplasm and forming unique ’inclusion-bodies.’ Once free in the cytoplasm, they also invariably develop dense and uniform coatings of cytoplasmic ribosomes on their surfaces, and finally, they become surrounded by ’annulate’ lamellae’ of the ER. In the end, these annulate-lamellar enclosures become the ultrastructural ’signatures’ of these inclusion-bodies, and serve to positively and definitively identify all cells that have been effectively transfected. Importantly, these new EM-observations define several new and unique properties of these classical Lipofectamines, and allow them to be discriminated from other lipoidal or particulate transfection-reagents, which we find do not physically break out of endosomes or end up in inclusion bodies, and in fact, provoke absolutely none of these ’signature’ cytoplasmic reactions.

## INTRODUCTION

The goal of this study was to describe the morphological features associated with the entry of the commonly-used transfection-reagents into cells, and the delivery of their DNA or RNA- payloads into cells, using primarily classical thin-section electron microscopy (EM) of fixed cells, but also using our unique form of ’quick-freeze, deep-etch’ EM, where possible. There have been a number of previous efforts to do this over the decades since the introduction of transfection-reagents, but besides concluding that they enter cells by endocytosis, and not by direct penetration through the plasmalemma (PM), relatively little has been learned.

Because the cationic lipoplexes that carry the plasmids or polynucleotides in or on them are generally globular structures, ranging in size up to nearly a micron or so, they are too big to fit into clathrin coated endocytotic pits on the cell surface. Their mechanism of uptake has always been seen as some form of macropinocytosis, either active - by protrusion of cellular extensions - or passive *via* the cationic lipids sticking to the PM tightly enough to ’wrap’ themselves in it, until they form a closed compartment (or perhaps a combination of these two mechanisms). (Colin et al., 2000; Friend et al., 1996; Hanson et al., 2010; Le Bihan et al., 2011; Li et al., 2019; Lin et al., 2014; Malatesta, 2021; Plaza-Ga et al., 2019; Reifarth et al., 2018; Verma et al., 2008; Zabner et al., 1995; Zhou and Huang, 1994).

All previous studies have agreed, though, that lipoplexes end up in relatively large or strikingly large endosomes, which are presumed by all to interact, as usual, with preexisting endosomal structures and/or lysosomes. and soon thereafter become acidic inside as they acquire proton pumps during the usual maturation of endosomes. This presumably protonates their DNA or RNA and allows it to dissociate from the cationic lipoplex-carrier, and somehow then to enter the cytoplasm. (Caracciolo et al., 2009; Colin et al., 2000; Hoekstra et al., 2007; Lazebnik et al., 2016; Le Bihan et al., 2011; Liu et al., 2004; Magalhaes et al., 2014; Midoux et al., 2009; Singh et al., 2004; Varkouhi et al., 2011). However, no thin-section EM-study to date has managed to directly demonstrate this entry, nor to elaborate upon what happens to these ’foreign’ macropinosomes loaded with lipoplexes, over longer residence-times in the cell.

In this regard, we were struck by certain unusual cytoplasmic bodies that had been observed in two EM-studies of transfected cells, which consisted of spherical ’bubbles’ uniformly surrounded by small electron-dense objects stuck to their surfaces (Asensio et al., 2013; Shu et al., 2011) (Figure 1, Panels A & B). The Shu et al paper, from the labs of Tsein and Ellisman at UCSD, was in fact the first introduction of the miniSOG technique (now evolved into APEX- labelling for EM), in which a light-activated peroxidase is conjugated to the protein of interest, and localized in the EM by DAB-staining, after the protein conjugate has been introduced into cells by transfection. Having transfected-in the gap-junction protein Connexin 43 fused to miniSOG, these authors described Fig1A as "black dots studding the membranes of trafficking vesicles, which may represent single gap-junction connexons." The Asensio et al paper, from the Edwards lab at UCSF, represented PC12 cells that had been transfected with miniSOG- VPS41, using Lipofectamine 2000, which they explained "resulted in about 25% of the transfected cells showing clustered membrane vesicles *decorated with regular, periodic arrays of electron-dense puncta*." They said no more about this, because they had already learned from us that we had begun to observe such structures in our own thin-section EM of cells exposed to Lipofectamine 2000 for entirely different reasons (Figure 1, Panel C). Namely, we were using Lipofectamine simply to enhance the cellular uptake of a prion-forming fibril- construct whose toxic effects we were studying with M. Diamond at the time (Boassa et al., 2013; Frost et al., 2009; Holmes et al., 2013; Kfoury et al., 2012; Kolay and Diamond, 2020; Luk et al., 2009; Sanders et al., 2014), a powerful technique developed by Nekooki-Machida et al., 2009. However, at about that time we were also observing the same sorts of cytoplasmic inclusions in a whole variety of cultures that we were transfecting with *anything and everything but* connexons. The inevitable conclusion was that Shu et al’s original description must have bee incorrect, and in the present study we will provide compelling evidence that the dark particulate bodies in question were not gap-junction connexons at all, but were attached ribosomes, fundamentally the same as the "ribosome-associated vesicles" or "RAVs" still being described in current studies of Lipofectamine 3000-transfected cells (Carter, Freyberg et al, 2020). For all these reasons, these earlier observations prompted us to undertake a systematic EM-evaluation of the mechanism of entry and the ultimate fate of all different sorts of transfection-reagents into cells, the results of which are reported here.

**Figure 1.**
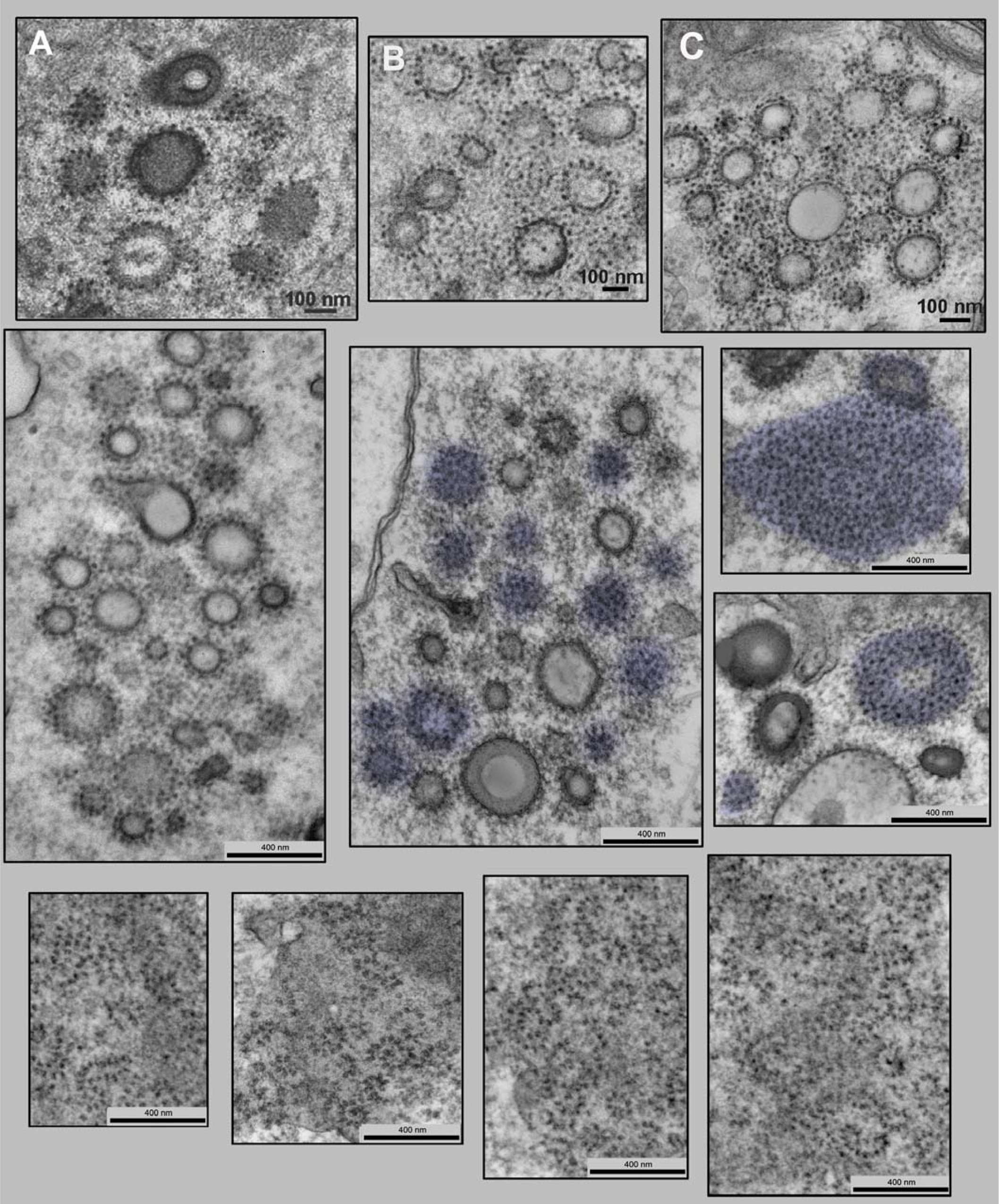
**<u>Panel A</u>:** Figure 4C from the 2011 paper by Shu, Deerinck, Ramko, Davidson, Jin, Ellisman and Tsien (Shu et al., 2011) which they explained "represented HEK293 cells that had been transfected 24 hours earlier with a plasmid containing the gap-junction protein Connexin 43 fused to a miniSOG". They explained that this resultant EM showed "black dots studding the membranes of trafficking vesicles (center-to-center distance ∼14 nm)", which they concluded "*may represent single gap-junction <u>connexons</u>*." **<u>Panel B</u>:** Supplemental Figure 4 from the 2013 paper by Asensio, Sirkis, Maas, Egami, Brodsky, Shu, Cheng and Edwards (Asensio et al., 2013), which they explained "represented PC12 cells that were transfected with VPS41 siRNA and RNAi-resistant miniSOG-VPS41, using Lipofectamine 2000 according to the manufacturer’s instructions." They explained that "this resulted in about 25% of the transfected cells showing clustered membrane vesicles *decorated with regular, periodic arrays of electron-dense puncta*." They elaborated on this no further, because by then we had made them aware of our results, shown next. **<u>Panel C</u>:** Our own thin-section EM of a cell exposed to Lipofectamine 2000 not for transfection, but for an entirely different reason: to enhance the uptake of a prion-forming fibril-construct whose toxic effects we were studying with Mark Diamond at Washington University circa 2013, a technique developed by Nekooki-Machida et al., 2009. (See Boassa et al., 2013; Frost et al., 2009; Holmes et al., 2013; Kfoury et al., 2012; Kolay and Diamond, 2020; Luk et al., 2009; Sanders et al., 2014.) Clearly, the structural-parallels with Panels A and B are obvious. **<u>Middle Panels:</u>** Examples of the characteristic clusters of multilaminar spherical bodies that invariably accumulate in Lipofectamine 2000-transfected cells. These ’inclusions’ generally exclude most other cytoplasmic organelles, but invariably display coatings of ∼20nm diameter dense particles all over their surfaces, which we presume are ribosomes. Where the bodies are thin-sectioned in a manner that displays their surfaces *en face,* they are highlighted in blue. This demonstrates that no matter how densely and uniformly they become coated with the small dense particles, these presumptive ribosomes remain strictly *singular*. They never form any complexes like polysomes. Instead, they space themselves almost perfectly uniformly all over each spheroid, and remain separated from each other by some 15-30nm, thereby assuming almost perfect trigonal symmetry. Scale bars, all 400nm. **<u>Lower panels</u>:** Contrasting dramatically with the uniform spacing and symmetry of the presumptive ribosomes on the Lipofectamine 2000 inclusions above, four fields of rough endoplasmic reticulum (RER) are chosen to show at the same magnification the characteristic appearance of the curvy polysomes that normally attach to RER. Clearly, when ribosomes are aligned into polysomes along mRNA, they approximate each other much more closely than they do on the surfaces of the Lipofectamine-inclusions, above. Scale bars, all 400nm.

It is important to stress at the outset that in a way we have been ’working in the dark’ here, because the detailed chemical composition and formulation of most *commercial* transfection reagents are trade-secrets which we are not permitted to know (Dalby et al., 2004; Hawley-Nelson and Ciccarone, 2001; Hawley-Nelson and Ciccarone, 2003; Hawley-Nelson et al., 2008; Ohki et al., 2001). However, since one of the authors of this paper (P. Felgner) is the actual inventor of the original transfection-reagents upon which all subsequent commercial reagents have been based (Duzgunes and Felgner, 1993; Duzgunes et al., 1989; Felgner et al., 1994; Felgner, 1997; Felgner, 1999; Felgner et al., 1987; Konopka et al., 1997; Konopka et al., 1996; Malone et al., 1989; Stephan et al., 1996), we were fortunate enough to be able to compare our results with the various *commercial* reagents we could obtain, with results using better-defined cationic-lipoplexes manufactured by the Felgner lab, and we could thereby keep ourselves ’on track’.

The fundamental conclusion of this study is that we <u>still</u> do not know what happens to most transfection-reagents after they get endocytosed and they come to reside - seemingly indefinitely - in their own unique form of endosome; but for a few, and especially for the most commercially successful ones called by the trade names of Lipofectamine 2000 and 3000, we have learned that, in fact, they break out of their endosomes and come to lie free in the cytoplasm, in a unique form of ’inclusion body,’ just like those that were glimpsed in the handful of aforementioned studies (Asensio et al., 2013; Shu et al., 2011, Carter, Freyberg et al, 2020)(Figures 1A & B).

The structural signature of this breakout, and the details of the unique and pathognomonic cellular response that it provokes in the endoplasmic reticulum (ER) of the cell, are the subjects of this study. But again, we should stress that this whole sequence happens, as far as we have seen so far, with only the transfection reagents called Lipofectamine 2000 and 3000, and not with other Lipofectamines, nor with the classical lab-produced ’lipofectins’, nor with a number of other commercial transfection reagents that we have tested. It seems to be unique to these two reagents. This may eventually help to explain the particular <u>effectiveness</u> of these two reagents, which has led to their wide adaptation in the field. But because the observed cellular reaction described here appears to occur only with these few, it remains to be seen whether it is an obligate part of the injection of polynucleotides into cells by transfection reagents, in general, or is simply an epiphenomenon that helps to explain something about the potential cytotoxicity of these particular reagents, when applied in excess (Ballarin-Gonzalez and Howard, 2012; Basha et al., 2011; Bauer et al., 2006; Damodaran et al., 2020; Dokka et al., 2000; Filion and Phillips, 1997; Konopka et al., 1996; Li et al., 2019; Lozhkin et al., 2017; Lv et al., 2006; Mo et al., 2012; Napoli et al., 2017; Niso-Santano et al., 2011; Terada et al., 2021b).

## RESULTS

### 1. Description of the Lipofectamine inclusion-body

Before describing the method of entry of the Lipofectamines, or describing their initial endocytic residences, or the unique and characteristic ER-reaction as they break out of their endosomes, we first present here their most striking and characteristic configuration, once they are free in the cytoplasm as seen in HEK293 cells. These are micron-sized clusters of dozens of multilamellar spheroidal-bodies that exclude all other cytoplasmic organelles, except for a characteristic monolayer of ∼20nm diameter dense particles that accumulate uniformly all over their surfaces (Fig.1, middle panels).

These tiny dense particles we presume are ribosomes, likely to be attached electrostatically to the surfaces of the cationic lipid droplets, due to their own strong negative surface-charge, which has been well documented (Baker et al., 2001; Bauer and Traub, 1995; Blad et al., 1977; Blad-Holmberg, 1979; Knight et al., 2013; Schavemaker et al., 2017; Traub et al., 1998; Trylska et al., 2004). Further reason to believe that these tiny dense bodes are ribosomes is not only that they are the same size as ribosomes, but they stain exactly the way that ribosomes do. Specifically, their reactivity with uranyl acetate block-stains and grid-stains match all the classical descriptions of ribosomal-staining (Afzelius, 1992; Bawdon et al., 1984; Beiras et al., 1987; Brodie et al., 1982; Dvorak and Morgan, 2001; Kislev et al., 1965; Korn et al., 1985; Morgenstern, 1977; Payne et al., 1983; Piefke et al., 1986; Sawaguchi et al., 2001; Soloff, 1973; Voigt et al., 2002). Likewise, their reactivity with lead-stains exactly matches all the classical descriptions of ribosomal staining (Bernhard, 1969; Brodie et al., 1982; Dallner et al., 1966; Farquhar and Palade, 1965; Huxley and Zubay, 1961; Karnovsky, 1961; Reynolds, 1963; Venable and Coggeshall, 1965).

Despite how densely and uniformly the Lipofectamine-spheroids become coated with ribosomes, these ribosomes remain strictly *singular*. They never form any complexes like polysomes (cf. Fig 1, lower panels), but instead, space themselves absolutely uniformly all over each spheroidal body, thus displaying in the EM near-perfect trigonal symmetry. This is particularly apparent whenever these spheroidal bodies happen to be viewed in glancing thin-sections, which display their surfaces *en face* (Fig.1, middle panels). This uniform and characteristic spacing contrasts dramatically with how closely ribosomes approximate to each other when they aligned into polysomes along mRNA, as they are usually observed in the general cytoplasmic milieu of the cell, or when attached to the rough endoplasmic reticulum (Fig 1, lower panels). Most importantly, this characteristic ribosome-decoration is <u>not</u> seen when the Lipofectamine droplets are still inside of endosomes (Fig.2), or when their endosomes are first breaking down and provoking the characteristic ER-reaction, as will be described and illustrated below.

**Figure 2.**
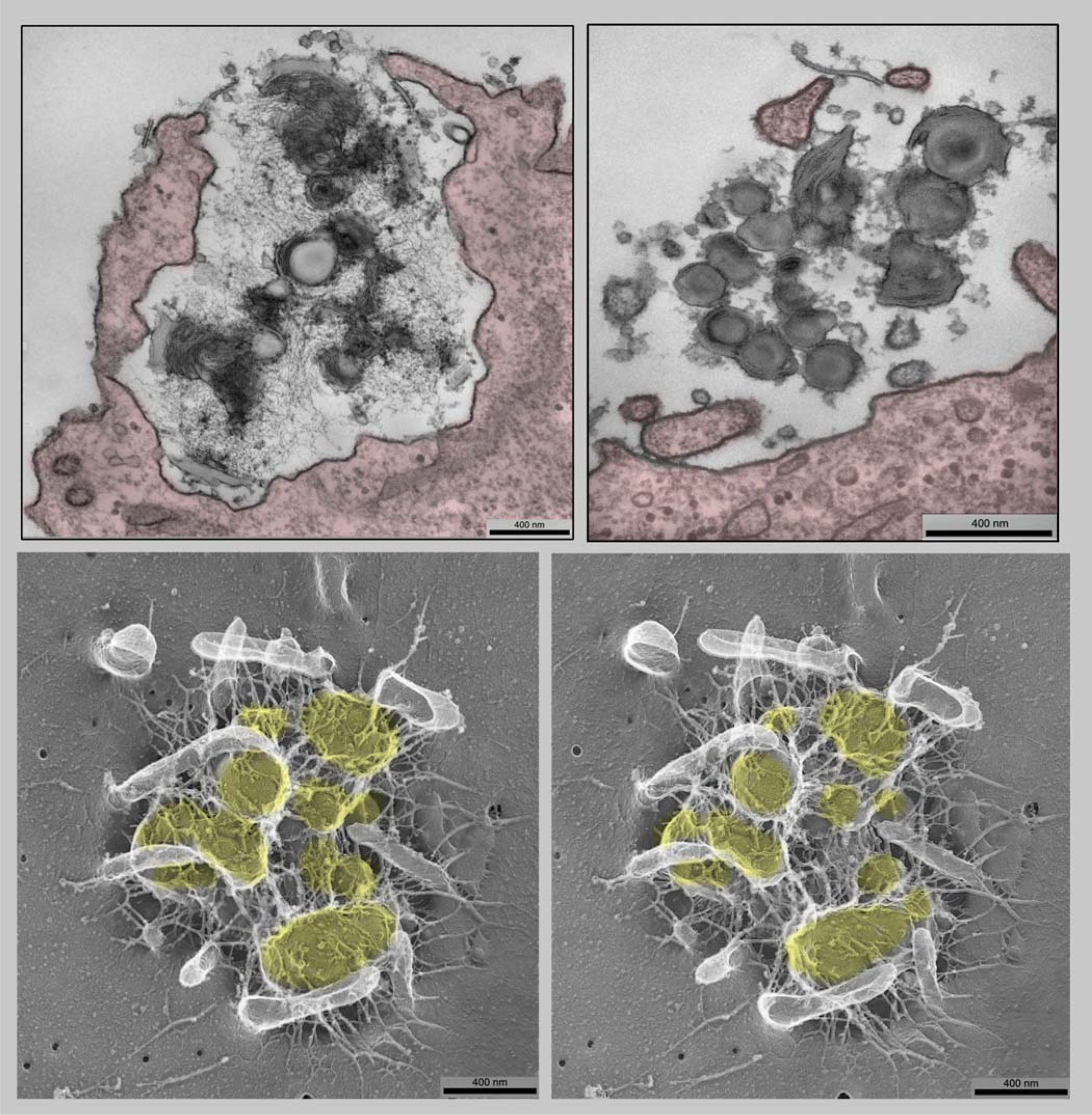
**Upper left panel:** The appearance of Lipofectamine 2000 -DNA complexes outside of cells, and attached to their surfaces in large invaginated domains. (Presumably, these are developing macropinosomes.) These consist of micron-sized clusters of multilamellar spheroids, individually some 100-200nm in diameter, which are invariably coated with thin ’hairs’ that presumably represent the attached plasmid DNA. (We find this appearance to be completely consistent, regardless of how differently our collaborators have prepared their particular Lipofectamine-DNA mixtures.) Note that some of the multilamellar spheroids look broken or incomplete, as if they had ’cracked apart’. This appearance will persist throughout their lifetime in the cell, even as they break out of their macropinosomes into the cytoplasm. Scale bar 400nm **Upper right panel:** In contrast, the appearance of <u>naked</u> Lipofectamine 2000 attached to the cell surface (that is, Lipofectamine *all by itself*, without ever having been complexed with any DNA). It again consists of micron-sized clusters of multilamellar spheroids and fragments of spheroids; but importantly, the coating of thin ’hairs’ seen to the left, which presumably represents the plasmid DNA, is <u>completely absent</u> from such views. Instead, the spheroids display here and there on their surfaces irregular particles and the occasional exosome, which we presume have derived from scattered debris in the culture medium, since this ’decoration’ is not seen when we image Lipofectamine all by itself straight from the bottle. Scale bar 400nm **Lower panels:** Stereo-view of the "deep-etch" EM-appearance of a Lipofectamine 2000+DNA complex when it is attached to the surface of a cell, in a shallow indentation presumably representing incipient macropinocytosis, prepared by our traditional ’deep-etch’ EM-technique (Heuser & Salpeter, 1979; Heuser 2000a,b). As in the left panel above, the complex consists of a micron-sized cluster of roughly spherical bodies, individually some 200-400nm in diameter (highlighted yellow here), again invariably coated with exceedingly thin ’hairs’ that presumably represent their plasmid DNA. Such "deep-etch" EM’s graphically display that these ’hairs’ are tangled and branched, but quite uniform in caliber (∼8nm). Clearly, they have a strong affinity for the plasmalemma, and appear to tether the spherical Lipofectamine bodies to the cell surface. Likely as part of the macropinocytotic process, the cell itself has erected slender microvilli (white in this contrast-reversal of the original platinum replica of this freeze-dried cell), which look like they are trying to ’embrace’ the transfection-complex. Scale bars, all 400nm.

One important consequence of this unique ribosome-decoration is that the Lipofectamine spheroids or broken spheroids no longer contact each other, as they do in their initial encounter with the outsides of cells, or when they reside in endosomes, or are in the process of breaking out of the endosomes in which they come to reside (described below). Moreover, our EM’s demonstrate clearly that these ribosome-studded clusters of lipofectamine-droplets remain in well-defined and partitioned domains within the cytoplasm, domains that utterly exclude all other cytoplasmic organelles, which can therefore only be considered to be separate inclusion-bodies or ’condensates’ in the cytoplasm (Alberti and Hyman, 2021; Banani et al., 2017; Brangwynne et al., 2009; Hyman and Brangwynne, 2011; Hyman et al., 2014; Wheeler and Hyman, 2018; Woodruff et al., 2018). Furthermore, they continue to remain in such domains, even when the whole cytoplasm becomes compacted during apoptosis, which is the fate of a large proportion of Lipofectamine-transfected cells (cf., Supplementary. Fig.1).

Importantly, we stress again that it is clearly apparent from all our EM’s that such decoration with ribosomes does not occur until the Lipofectamine droplets are fully released from their incoming endosomes and are fully released from the protective coatings of ER that form initially around these endosomes as they begin to rupture inside the cell, as will be shown below.

### 2. Initial appearance of the Lipofectamines as they enter cells

Regardless of which laboratory around the world provided us with transfected cells, and regardless of the details of how they prepared their particular Lipofectamine-DNA mixtures, the appearance of Lipofectamine when it is <u>outside</u> of cells, and attached to the surface of cells, has been completely consistent. It appears as micron-sized tight clusters of multilamellar spheroids, individually some 100-200nm in diameter, invariably coated with thin ’hairs’ that presumably represent the plasmid DNA they are intended to import (Fig.2, left panel). Our complementary "deep-etch" three-dimensional stereo-EM’s of these clusters (Fig.2, bottom panels) demonstrate that these ’hairs’ are quite uniform in caliber, and if we subtract from their apparent thickness the 3nm of platinum that we apply to image them in such freeze-dried preparations, we find that they are only a few nanometers in width, as would be expected for the DNA itself. (See Supplemental Figure 2, for examples of how such stereo-EM’s can be converted into 3-D ’anaglyphs,’ for ease in viewing the hairy DNA that attaches to these lipoplexes (Heuser 2000a,b).) In every respect, these resemble the ’hairs’ seen on the surfaces of cationic lipoplexes in all the early freeze-fracture studies of them (Aytar et al., 2012; Donkuru et al., 2010; Dunlap et al., 1997; Sternberg et al., 1998; Sternberg et al., 1994; Xu et al., 1999); but to our eye, they do not look *thick enough* for the DNA to be coated with its own monolayer or bilayer of lipid, as most of the earlier structural studies concluded (Duzgunes and Felgner, 1993; Duzgunes et al., 1989; Felgner et al., 1994; Felgner, 1997; Felgner et al., 1987; Konopka et al., 1996; Malone et al., 1989; Stephan et al., 1996). We tentatively conclude that they are the naked DNA, itself.

These peripheral ’hairs’ are not only readily apparent in all our deep-etch and thin-section EM’s of Lipofectamine-DNA attached to the outsides of cells (Fig.2), but most importantly, we find that they are <u>utterly absent</u> when we apply Lipofectamine *all by itself*, without ever complexing it with any DNA before exposure to cells (Fig.2, right panel). (It is important to add here that the tight clusters of multilamellar spheroids attached to the cells without DNA look otherwise identical to the ones that <u>do</u> have DNA) (cf., compare right and left panels in Fig.2).

It is important to note here also that in all our EM’s, the liberated lipoplex-spheroids themselves look either uniform and amorphous inside, or else, more commonly, look multi-laminated inside. This fits with most of the early freeze-fracture studies of such complexes, which also concluded that they are multilamellar (Aytar et al., 2012; Donkuru et al., 2010; Dunlap et al., 1997; Sternberg et al., 1998; Sternberg et al., 1994; Xu et al., 1999), but it specifically and strikingly disagrees with most of the subsequent cryo-EM that has been done on lipoplexes (Battersby et al., 1998; Ewert et al., 2021; Fenske and Cullis, 2005; Gustafsson et al., 1995; Huebner et al., 1999; Kulkarni et al., 2018; Kulkarni et al., 2020; Kulkarni et al., 2019a; Kulkarni et al., 2019b; Kuntsche et al., 2011; Leung et al., 2015; Maurer et al., 1999; Radler et al., 1997; Ramezanpour et al., 2019; Safinya et al., 2014; Silva et al., 2014; Terada et al., 2021a; Wan et al., 2014). Clearly, this is a fundamental discrepancy that must be dealt with in the future, and when considering all the models of how these lipoplexes are actually constructed (Ewert et al., 2005; Koltover et al., 1998; Leal et al., 2010; Lin et al., 2003; Majzoub et al., 2015; Majzoub et al., 2016a; Majzoub et al., 2016b; Radler et al., 1997; Safinya et al., 2014). But we would point out that the methods of generating such lipoplexes have evolved ofer the years, and currently involve much more modern techniques of microfluidics than were used in the past, when the Lipofectamines were first introduced.

Parenthetically, it is important to note here that our experiments with Lipofectamine that has no DNA or RNA associated display the same patterns of entry and of eventual decoration with ribosomes as lipofectamine <u>with</u> its full complement of plasmid DNA, so it cannot be the DNA, itself, that causes the endosomal rupture, the ER-recruitment, or the eventual ribosome- decoration of the spheroids that we consistently observe. (It is also important to note that no such thin fibrils or ’hairs’ are <u>ever</u> seen to remain on any of the ribosome-decorated spheroids, once they come to reside in the cytoplasm; so the DNA must come off of them by the time they are freed into the cytoplasm.)

While residing within endosomes, the Lipofectamine-spheroids become so tightly compacted that it is impossible to determine, either by freeze-fracture EM or by thin-section EM, whether they retain their surface-coating of DNA (Fig.3); but we presume, from the images to be presented next, that eventually the acidic interior of the endosome protonates their polynucleotides and separates them into free DNA and concentrated phospholipid droplets - - droplets that are sufficiently hydrophobic to attach and weaken the endosomal membrane and eventually rupture it, as described next.

**Figure 3.**
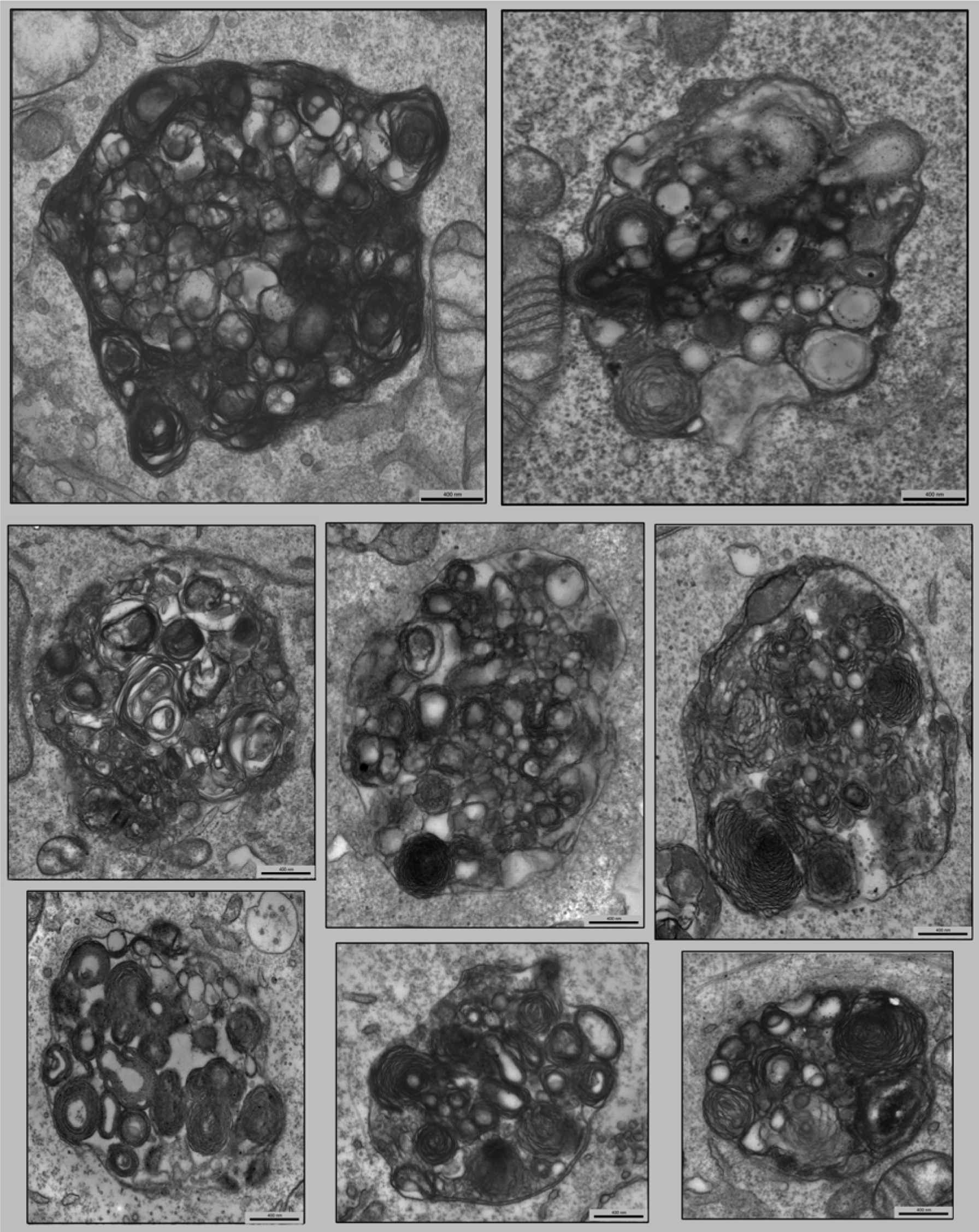
Examples of the sorts of endosomes that are *<u>never</u>* seen in untransfected cells, and are presumed to be macropinosomes that are chock-full of Lipofectamine 2000. While residing within such unusual endosomes, the multilamellar-spheroids become so tightly compacted that it becomes impossible to determine, either by ’deep-etch’ EM or by thin-section EMs such as these, whether they still retain their surface-coating of DNA. Scale bars, all 400nm.

### 3. Endosome rupture with endoplasmic reticulum (ER) recruitment

The most intriguing and elusive step in this transmission of Lipofectamine from outside of the cell to inside the cell is the stage of endosome-rupture. Naturally, this must be a fleeting process, akin to vesicle exocytosis, which has been notoriously difficult to capture over the decades, without heroic efforts to coordinate it with fixation or with cryo-immobilization of the cell (Heuser, 1981, 1983, 1989a,b; Heuser, 2011; Heuser et al., 1979). But there is no way known, still to this day, to coordinate endosome-rupture with anything like this precision.

Consequently, the observer must search through vast numbers of cells, in hundreds of transfected cultures, to obtain images that suggest anything like endosome-rupture, or to capture its near-aftermath.

Here, in attempting to ’catch’ the moment of the entry of Lipofectamines into the cell *via* endosome-rupture, we were greatly aided by our discovery, serendipitously, of another cellular reaction that appears to precede or accompany it. This is a cellular reaction that involves a wholesale re-arrangement of elements of the endoplasmic reticulum (ER), and which is readily visible and recognizable in the EM, and which presumably takes quite some time to develop, and then takes quite some time to resolve after the endosome-rupture. Specifically, this re- arrangement involves the formation of self-enclosed multilayers of ER, connected to one another by narrow channels, which no doubt form formidable barriers to egress of the newly-exposed Lipofectamine-spherules into the cytoplasm proper (Fig.4).

**Figure 4.**
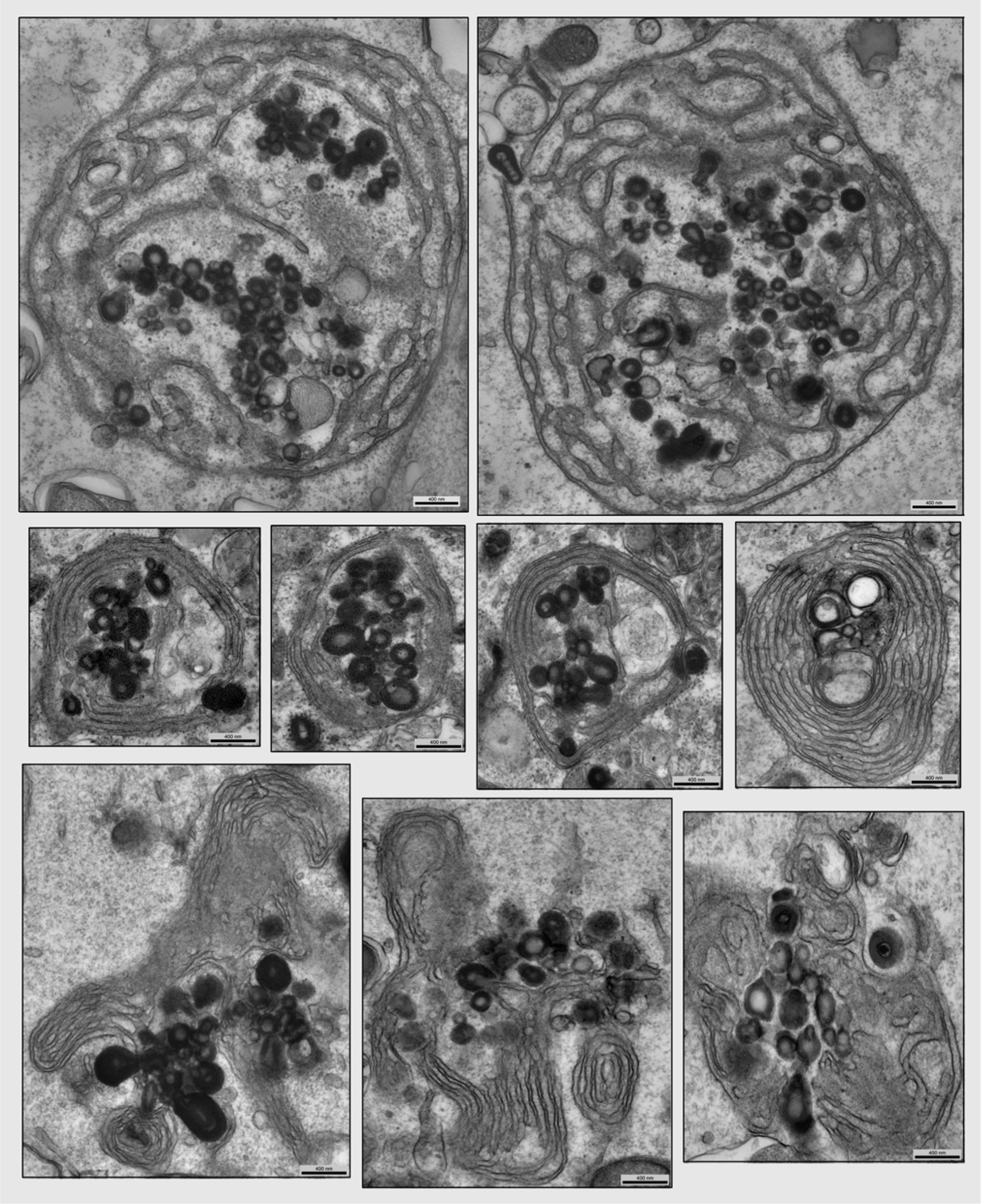
**Middle four panels:** Typical examples of the cellular reaction that accompanies the incipient rupture of Lipofectamine 2000-containing macropinosomes. This involves a wholesale re- arrangement of elements of the endoplasmic reticulum (ER) into self-enclosed multilayers, connected to one another by narrow channels, which are presumably erected to form a barrier to the egress and escape of the Lipofectamine-spherules from the endosomes into the cytoplasm, proper. However, it is apparent from these views that the spherules have already begun to escape into the cytoplasm anyway, although they have not (yet) acquired the characteristic coatings of ribosomes seen in Fig.1. Scale bars 400nm. **Upper two panels:** Two more examples of the cellular reaction that accompanies the rupture of Lipofectamine 2000-containing macropinosomes, representing we believe a slightly earlier stage than below. At this point, the re-arrangement of elements of the endoplasmic reticulum (ER) into self-enclosed multilayers is ’looser’, and thus the narrow channels interconnecting the layers are more apparent. Clearly, the cytoplasm within these multilayer-enclosures looks relatively pale, partly because it not appear to harbor any free ribosomes. Even if these multilayers were erected to form a barrier to incipient endosome rupture, it is apparent from these views that the spherules have escaped into the cytoplasm anyway; although again, they have not (yet) acquired the characteristic coatings of ribosomes seen in Figure 1. Scale bars 400nm. **Lower three panels:** Illustration of the remarkable persistence of the ER-elaborations displayed above. Such tangled multilayers of endoplasmic reticulum persist around the peripheries of the Lipofectamine inclusions, *long after* the inclusions have consolidated, and their spherules have all become uniformly decorated with ribosomes. Indeed, they remain abundantly observable, apparently clinging to the edges of the Lipofectamine/ribosome agglomerations, throughout the whole duration of transfection. Scale bars 400nm.

The fundamental composition of the cytoplasm within these multilayer-enclosures is visibly different from the surrounding cytoplasm. In thin sections, it appears extremely pale, as if it were depleted of the normal complement of soluble cytoplasmic components (Fig.4, upper panels). Moreover, most importantly - - and critical to the scenario being elucidated here - - this milieu contains *absolutely no ribosomes*. And equally critically, the Lipofectamine spherules released from the broken endosomes into this pale environment are not yet decorated <u>at all</u> with any ribosomes (see again Fig.4, upper panels). This single ultrastructural feature strongly suggests that the elaborated ER around them is ’protecting’ or ’enclosing’ the freshly-released Lipofectamine, or at least is *attempting* to do so. In this and many other ways, this ER- elaboration reminds us of early forms of autophagocytosis (Biazik et al., 2017; Biazik et al., 2015; Fengsrud et al., 1995; Gudmundsson et al., 2022; Hayashi-Nishino et al., 2009; Kovacs et al., 1982; Park et al., 2020; Yla-Anttila et al., 2009; Zwilling and Reggiori, 2022) and strongly suggest that the cell is attempting to encapsulate or ’insulate’ itself from the Lipofectamine- droplets that are emerging from the breaking endosomes. That this ’insulation’ is partially successful is most notably indicated by the absence of any ribosome-decoration of the Lipofectamine, at this early stage (cf., Fig.4, upper panels).

Also remarkable in all the thin-sections of Lipofectamine-transfected cells is the remarkable persistence of these ER-elaborations, these tangled multilayers of endoplasmic reticulum, <u>long</u> <u>after</u> they are formed. They remain commonly and abundantly observable, apparently clinging to the edges of the Lipofectamine/ribosome agglomerations, throughout the whole duration of transfection (Fig.4, lower panels). Even until the stage of apoptosis, remnants of this ER- reaction can often be observed, wherein they are *still* typically found immediately adjacent to the condensed Lipofectamine-aggregates (see again Supplementary Fig.1).

However, we should also stress that despite the obvious extensiveness and persistence of these ER-elaborations, we have never observed, within them, the formation of any of the classical "isolation membranes" that structurally define autophagocytosis (Biazik et al., 2017; Biazik et al., 2015; Fengsrud et al., 1995; Gudmundsson et al., 2022; Hayashi-Nishino et al., 2009; Kovacs et al., 1982; Park et al., 2020; Yla-Anttila et al., 2009; Zwilling and Reggiori, 2022). So, despite the obvious impression that the ER is attempting to ’wall off’ or ’shore up’ the breaking endosomes, the lipoplexes that are thereby released into the cytoplasm must not recruit LC3 or activate any of the rest of the normal cellular autophagocytosis-program.

Another ultrastructural characteristic of this ER-elaboration that may eventually help to explain what is happening at this point is that the subjacent laminae of ER in the tangled multilayers often appear in thin sections to be ’glued together’ by cytoplasm that is considerably more electron-dense than normal (Fig.5). This closely resembles certain ER-multilayers that form during the unfolded protein response resulting from VAPB-mutations (Borgese et al., 2021; Fasana et al., 2010; Kanekura et al., 2006; Kanekura et al., 2009; Kuijpers et al., 2013; Navone et al., 2015; Nishimura et al., 2004; Papiani et al., 2012). More needs to be done, to see how and why these ER-changes are occurring in transfected cells, as well.

**Figure 5.**
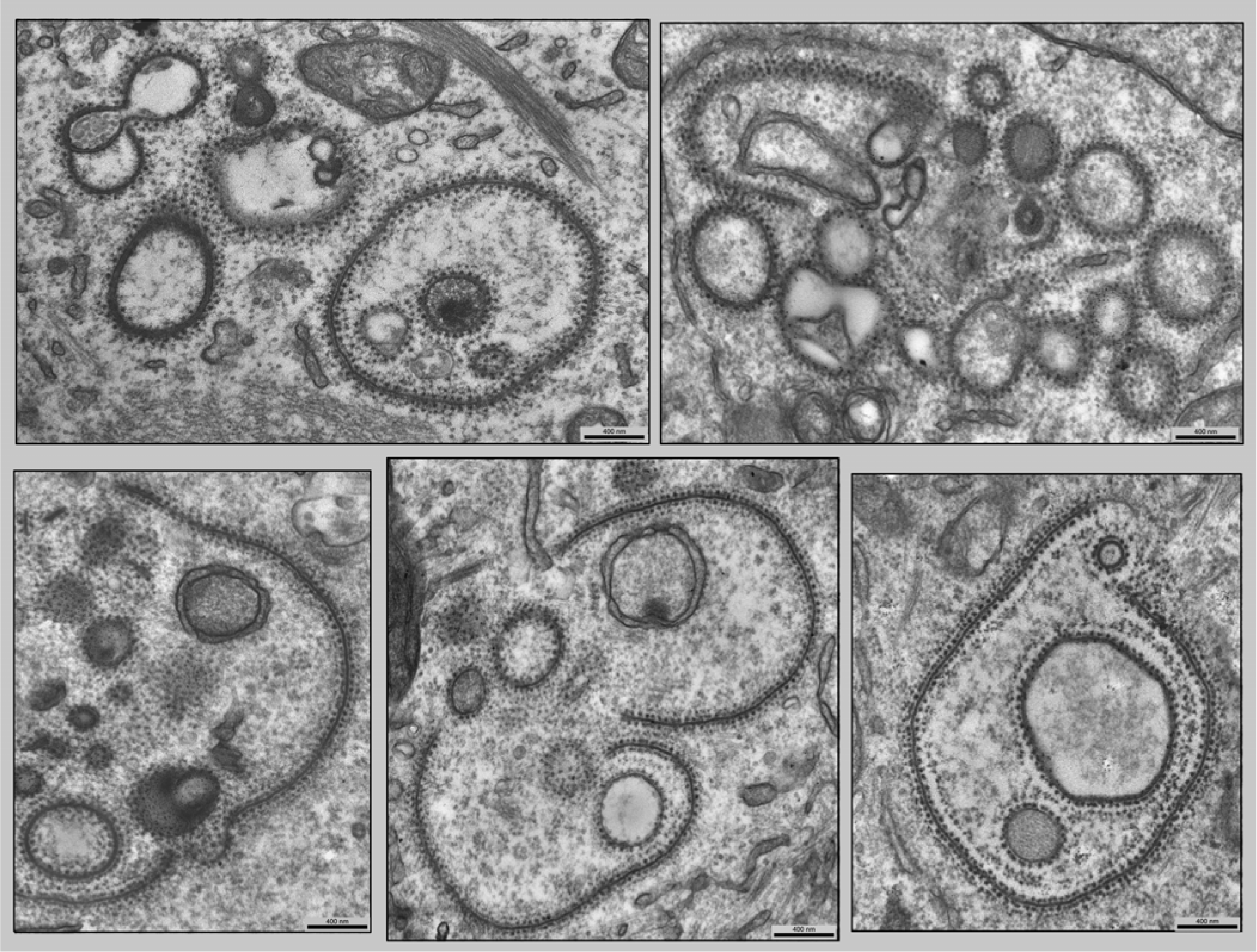
Examples of cells in which the endosomal membrane has clearly ruptured and released the lipofectamine droplets, and the few droplets that are visible have already become decorated with ribosomes, but where the *endosomal membrane itself* appears to have become decorated with ribosomes, suggesting that some transfer of cationic lipid to the endosome membrane *itself* must have happened during the endosome-rupture. Interestingly, the resultant ’broken- crescents’ of membrane are typically decorated with ribosomes *on both sides*, equally and uniformly. This dual-labeling on both sides of the thickened endosomal membrane demonstrates just how <u>regular</u> the spacing of ribosomes is under these conditions; it is nothing like the spacing of polysomes on RER (Fig.1). Scale bars 400nm.

### 4. Further ultrastructural changes that sometimes accompany the rupture of Lipofectamine-bearing endosomes

As mentioned above, ribosomes do not gain access to the lipid-droplets until the endosomal membrane *ruptures*. However, we often observe ribosome-decorated lipid drops that are surrounded by densly-staining fragments or crescents of membrane that have <u>themselves</u> become decorated with ribosomes (Fig.6). These could be interpreted as shredded or pulled- off layers of the Lipofectamine droplets *themselves*, or some other reconfiguration of them, which could concievably occur as they become exposed to cytoplasm. However, we prefer to interpret these ribosome-studded ’zippers’ of membrane as being fragments of the endosomal membranes *themselves*, left around after their rupture. If so, this would suggest that some sort of transfer of cationic lipids to the endosome membrane *itself* could be occurring, before or during its rupture, which would thereby give it, *also*, the characteristic ribosome-affinity described herein. Curiously, we observe that in many instances, these narrow ’broken- crescents’ of membrane are decorated with ribosomes *on both sides*, equally and uniformly (Fig.6). This makes it quite clear, due to the perfect symmetry and uniformity of the ribosome- decoration *on both sides of them*, that these crescent-shaped ’zippers’ must be biochemically the same on both their convex and concave faces, since both faces clearly have the same affinity for ribosomes. Clearly, more work needs to be done to fully understand these enigmatic (but characteristic) structures.

**Figure 6.**
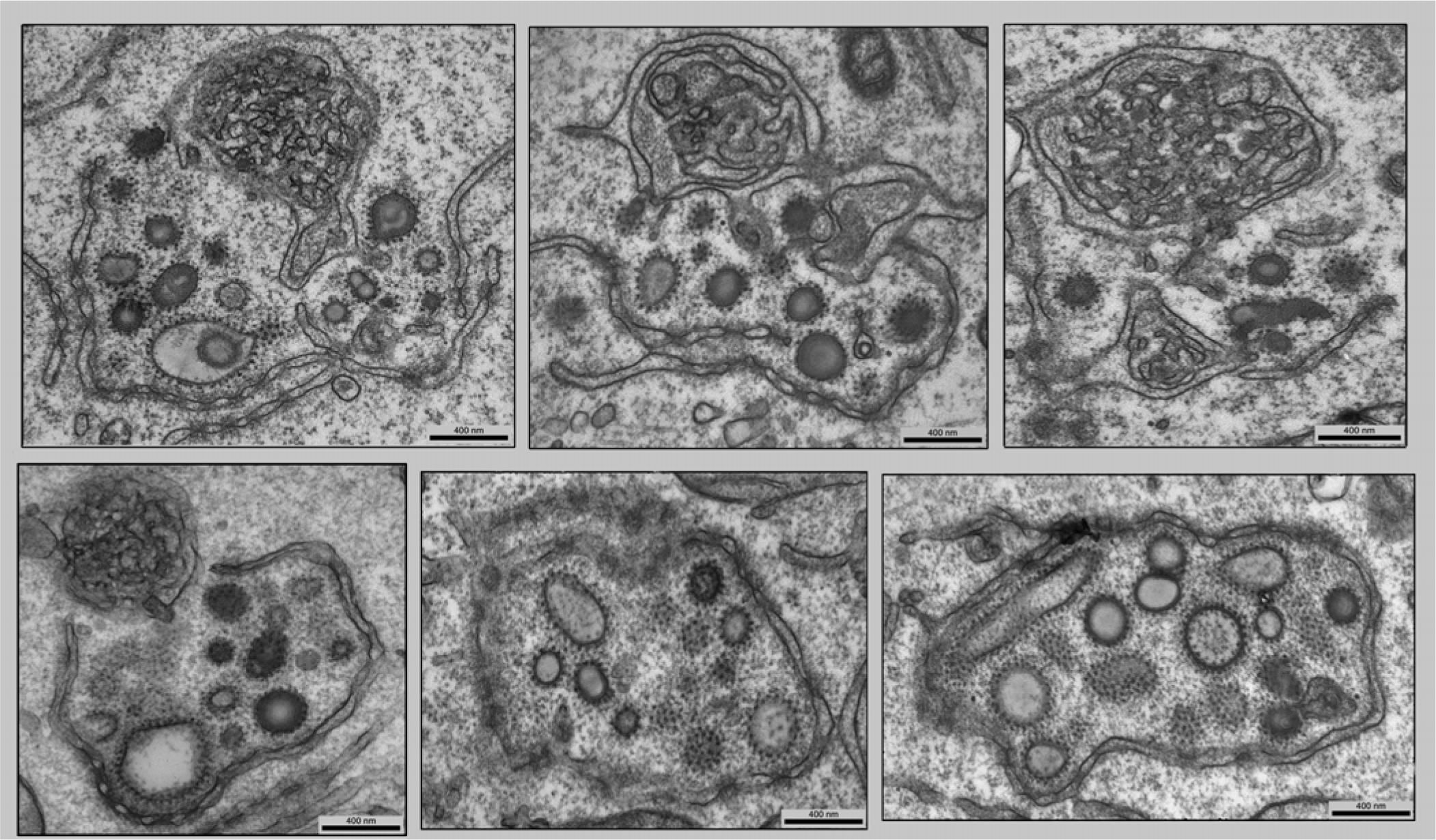
The final unique feature of all Lipofectamine-inclusions is that they end up enclosed in "annulate lamellae" (eg., nuclear-pore-studded domains of ER). These in fact define and delineate the outer-boundaries of the inclusions, themselves, and persist for the rest of the life of the transfected cell. Magnification 12,000x. Scale bars 400nm.

### 5. Final enclosure of the cytoplasmic Lipofectamine/ribosome phases by the endoplasmic reticulum’s "annulate lamellae"

Finally, another absolutely unique and notable ultrastructural feature that defines nearly all of the lipofectamine/ribosome inclusions that we have observed in the cytoplasm of transfected cells is that they finally end up enclosed in "annulate lamellae." These "annulate lamellae" are of course defined as nuclear-pore-studded domains of ER that occur in all cells, but are normally most elaborated in cells involved in rapid cell division or rapid nuclear envelope growth, such as oocytes or egg cells (Cordes et al., 1996; Erenpreisa et al., 2002; Eymieux et al., 2021; Haggag and Gilloteaux, 2005; Huber et al., 2020; Imreh and Hallberg, 2000; Kawabuchi et al., 1989; Kessel, 1983a; Kessel, 1983b; Raghunayakula et al., 2015; Walter and Tandler, 1989). In transfected cells however, we find that such annulate lamellae are invariably and strikingly over-abundant, and that they specifically direct themselves to enclosing the late Lipofectamine/ribosome inclusions, after the aforementioned ER- elaborations have retreated to their peripheries (Fig.7). There can be no confusion in differentiating these annulate lamellae from the aforementioned ER-elaborations that we concluded were trying to ’protect’ or ’contain’ the endosome-rupture. However, we never observed that <u>those</u> early ER-elaborations ever contained any nuclear pores, and commensurately, the annulate lamellae observed here never, ever, form tangled, interconnected multilayers. Hence, these must be two clearly distinct alterations of the cell’s normal ER.

**Figure 7.**
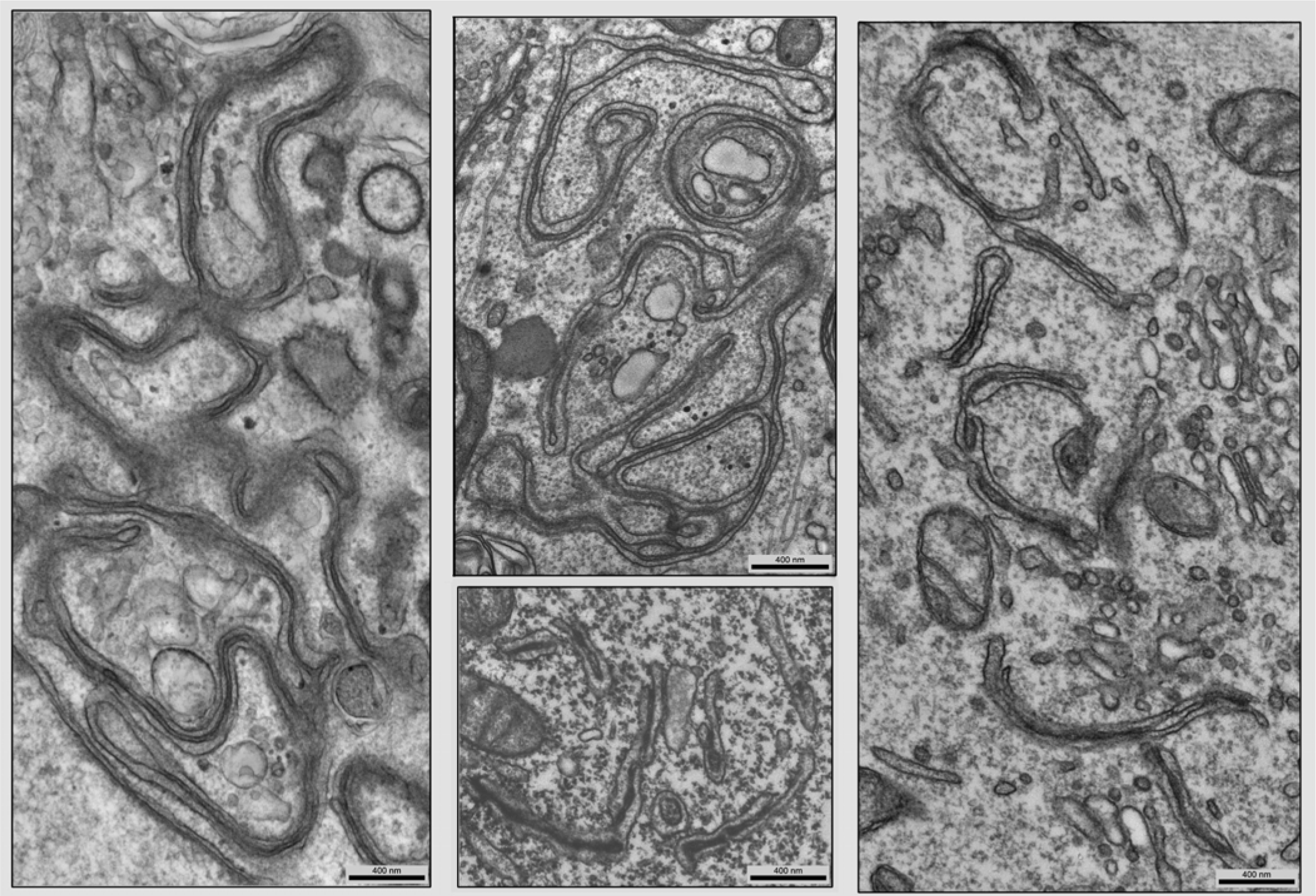
A further ultrastructural change that occurs in many but not all Lipofectamine 2000- transfected cells is that subjacent laminae of ER often appear in thin sections to become ’glued together’ by cytoplasm that is much more electron-dense than normal. In the text, we explain that these closely resemble the ER-multilayers that form in cells with ESCRT-mutations that mimic yeast cells’ "Class E" compartments, or in cells with VATB mutations that provoke the ER’s unfolded protein response (UPR). Why Lipofectamine provokes this reaction, and how it relates to the ER-changes seen in Figure 4, remains to be seen. Scale bars 400nm.

What ultimately recruits annulate lamellae to the cytoplasmic Lipofectamine inclusions remains a total mystery, but may turn out to have something to do with an affinity of the various positively-charged nucleoporin-proteins that are present in the outer cytoplasmic rings of all their nuclear pores, for all the ribosomes accumulated around the inclusions, themselves (Goryaynov and Yang, 2014; Lin and Hoelz, 2019; Mahamid et al., 2016; Unwin and Milligan, 1982). Still, our impression from years of witnessing transfected cells in the EM, is that annulate lamellae are generally and distinctly over-abundant in these cells *in general*…perhaps from defects or deficiencies in mitosis, from which all transfected cells are known to suffer (Brunner et al., 2000; Chernousova and Epple, 2017; Martin et al., 2014; Schiavone et al., 2000; Wang et al., 2018).

## DISCUSSION

As discribed in the Introduction, this entire study was initiated by observing, in cells exposed to Lipofectamine 2000, certain unique and characteristic cytoplasmic-inclusions that had been interpreted in earlier study to be sites of gap-junction connexon-synthesis, in mini-SOG transfected cells (Shu et al., 2011) (Figs.1A). Thereafter, we observed the same sorts of cytoplasmic inclusions in a whole variety of cultures that were exposed to Lipofectamine 2000 for a variety of other reasonss. Then, other groups began to describe the same sorts of cytoplasmic inclusions, in a whole variety of different sorts of transfections, but all using Lipofectamine or its congeners (Asensio et al., 2013)(Fig.1B); (Carter, Freyberg et al, 2020).

The conclusion was inevitable, that the original description of ’connexons" could not have been correct; and now, as documented in the present report, we realize that the dark particulate bodies in question were actually attached ribosomes.

Many questions are raised by the above observations. Most critical is the question of how we can be sure that they are actually ribosomes attached to the liberated Lipofectamine, and not some other sort of cytoplasmic dense-body. At the moment, we have only the staining-criteria that have been ’built into’ old-fashioned EM-lore: that ribosomes are osmiophilic, and that they stain characteristically with uranium and lead stains (Afzelius, 1992; Baker et al., 2001; Bauer and Traub, 1995; Bawdon et al., 1984; Beiras et al., 1987; Bernhard, 1969; Blad et al., 1977; Blad-Holmberg, 1979; Brodie et al., 1982; Dallner et al., 1966; Dvorak and Morgan, 2001; Farquhar and Palade, 1965; Huxley and Zubay, 1961; Kislev et al., 1965; Knight et al., 2013; Korn et al., 1985; Morgenstern, 1977; Payne et al., 1983; Piefke et al., 1986; Reynolds, 1963; Sawaguchi et al., 2001; Schavemaker et al., 2017; Soloff, 1973; Traub et al., 1998; Trylska et al., 2004; Venable and Coggeshall, 1965; Voigt et al., 2002). However, despite this strong evidence, we cannot be certain until we apply other modern light microscopic (LM) techniques, such as methods to stain the lipids in Lipofectamine and observe its spheroids directly, and then to immuno-stain these apparent ’ribosomes’ and image their staining in the LM, to thereby make a definitive CLEM-structural correlation.

Nevertheless, the conclusion is clearcut, that the most prominent cellular reaction to the uptake of the Lipofectamines into endosomes is the prompt attraction of elements of the endoplasmic reticulum (ER) to these particular endosomes. This ER invariably forms elaborate networks around the endosomes prior to or during their rupture, presumably in an attempt to ’contain’ or sequester or wall-off the released products. However, this attempt invariably fails, and the elaborated ER retreats to a slightly peripheral position, but a position that is still nearby the newly-generated hydrophobic inclusions (Fig.6). Additionally, besides this special and distinctive form of ER, the cell also recruits another form of ER, the form that is enriched in nascent nuclear pores (it’s so-called "annulate lamellae") to surround and enclose the Lipofectamine inclusions (Fig.7). In the end, these "annulate lamellar" compartments become the truly ’signature’-enclosures of the residual hydrophobic domains created by the released transfection-lipoplexes. This sequence of events is summarized diagramatically in Fig.8.

**Figure 8.**
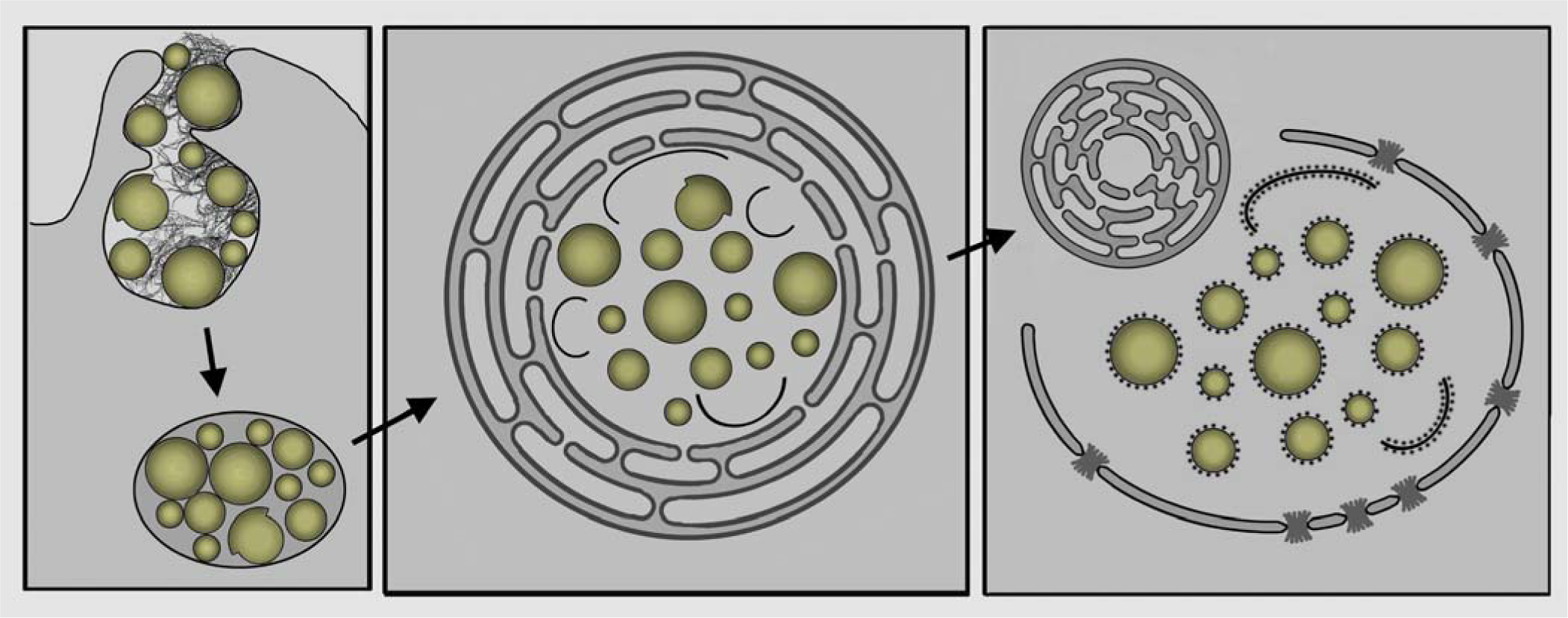
Diagram summarizing the cytoplasmic changes seen during Lipofectamine transfection. **Left panel:** Lipofectamine 2000 or 3000 droplets plus associated ’hairy’ plasmid DNA encountering cells gets endocytosed from the surface by macropinocytosis and sequestered in large macropinosomes, distinct from the rest of the endocytotic pathway. **Center panel:** Weakening and rupture of the macropinosome membrane provokes a dramatic reaction of the endoplasmic reticulum (ER), which surrounds and attempts to contain the Lipofectamine droplets that are becoming exposed directly to the cytoplasm. **Right panel:** Inevitably, this ER-containment fails and only remnants of it persist, while the freed Lipofectamine droplets and endosomal membrane-fragments become dramatically and distinctly decorated with cytoplasmic ribosomes, and the whole inclusion ultimately becomes surrounded by another form of ER that is rich in nuclear pores - - the so-called "annulate lamellae". Together, the structures seen here become the "signature’ of successful transfection, and by 24 hours, are found by EM in nearly <u>all</u> cells exposed to such Lipofectamine reagents.

Importantly, the EM-work described herein demonstrates that these ’signature’-enclosures clearly identify cells that have been successfully transfected. Namely, we consistently observe that their presence *correlates absolutely 100%* with whatever structural changes we were attempting to provoke by the particular genetic elements that we introduced by the transfection in the first place - - at least, it correlates 100% when these changes are successfully accomplished. However, we should also stress that we invariably find many cells in the EM that display <u>all</u> of the ’signature’-remnants of transfection described in this report, but that have obviously *failed* to properly express the genes that we originally attempted to inject by the transfection - - simply because they end up showing *none* of our intended structural alterations, but show <u>only</u> the aforementioned transfection-residua. This most likely indicates some failure in the plasmids we happened to use, themselves.

It is also very important to stress again that all the cytoplasmic changes described herein are <u>also</u> observed when the lipofectamines are applied *all by themselves*, without any DNA or RNA cargo at all. Moreover, it is important to stress again that in other collaborative work over the past two decades, we have also observed these same cytoplasmic changes whenever we use the Lipofectamines not as transfection-reagents, but as carriers for *non-nucleotide* cargos, like the prion-forming fibrils Tau and a-synuclein, which is a common procedure in the field of neuropathology (Boassa et al., 2013; Frost et al., 2009; Holmes et al., 2013; Kfoury et al., 2012; Kolay and Diamond, 2020; Luk et al., 2009; Nekooki-Machida et al., 2009; Sanders et al., 2014).

Hopefully, this primary report, which is clearly purely descriptive, will inspire further correlative LM-EM elaboration of the whole process of Lipofectamine transfection. Questions that are paramount to answer include (1) what happens to the DNA fibrils that associated with the outsides of the Lipofectamine-spheroids at the moment of uptake? (2) What proteins are involved in elaborating the convoluted ER that "attempts" to segregate and enclose the Lipofectamine as it is being released from endosomes into the cytoplasm? (3) What is the cell’s ’purpose’ in recruiting ribosomes to the liberated Lipofectamine, if any purpose at all? (4) What <u>else</u> is in the cytoplasmic inclusion that holds the Lipofectamine droplets together, in not-so-very-close proximity, but clearly excludes other cytoplasmic organelles, right up to the stage of apoptosis? What other proteins are there, and are these proteins (or RNAs) typical for other stress granules that have been studied? (Alberti and Hyman, 2021; Banani et al., 2017; Brangwynne et al., 2009; Hyman and Brangwynne, 2011; Hyman et al., 2014; Wheeler and Hyman, 2018; Woodruff et al., 2018). (5) Why does this ’filler’ in the inclusion attract annulate lamellae to its periphery? This has not been seen for any other cytoplasmic aggresome; not for stress granules, nor for P-bodies, nor for nuage in germ cells. Is <u>this</u> trying to tell us something about ribosome/nuclear pore interactions (Goryaynov and Yang, 2014; Lin and Hoelz, 2019; Mahamid et al., 2016; Unwin and Milligan, 1982)?

Finally, we must admit that even though purely EM-studies such as this one have always been demeaned by calling them "purely descriptive’, and even though we have always strived to overcome this criticism by establishing structure-function correlations with our EM whenever we could, this report suffers more than most, by our total ignorance of what is actually <u>inside</u> of Lipofectamine 2000 or 3000. Their composition and physical nature are closely-guarded trade-secrets that we have been unable to learn, but could be *critical* for explaining all the changes we have observed and reported here.

## MATERIALS AND METHODS

All the EM-images obtained in this study were from tissue cultures grown on glass-bottomed culture dishes, which allows for the cells to be imaged using LM for the structural changes actually intended by the particular transfection, as well as for a specific part of our EM-workup, as described below.

In general, HEK293A cells (Cat.R70507; Invitrogen) were maintained at 37°C and 5% CO_2_ in a humidified atmosphere and cultured in Dulbecco’s Modified Eagle Medium (DMEM-high glucose; Gibco, Life Technologies) containing 10% (vol/vol) FBS. Cell cultures were also supplemented with a 1% solution of penicillin/streptomycin (Gibco, Life Technologies) and 5 µg/ml of the prophylactic for all subsequent passages. For imaging studies, HEK293A cells were plated at a density of 300,000 cells/dish with a final volume of 1.5 mL on 29 mm circular glass-bottom culture dishes (#1.5; Cellvis) pre-coated with 0.01% poly-L-lysine solution (Sigma). The seeded cells were allowed to attach overnight prior to transfection with plasmid DNAs (0.1-0.2 μg/well) using Lipofectamine 2000 (2-5 μL/well; Invitrogen) within a small volume of Opti-MEM (200 μL; Invitrogen) according to the manufacturer’s instructions.

However, where indicated, long-term culture of cells overnight was accompanied by removing the media containing the Lipofectamine-complexed DNA at 4-6 h post-transfection and replacing it with complete culture medium. In general, cell densities were always kept between 50-80% confluence for the day of transfection to ensure maximal coverage of the culture surface and to facilitate the subsequent analysis by EM.

Except for the special use of glass-bottomed culture-dishes, the following protocol is the standard way that we prepare all samples for thin section EM (Heuser and Tenkova, 2020; Heuser, 2022). At the end of the transfection period, the tissue-culture medium was exchanged directly with our standard aldehyde fixative, which is 2% glutaraldehyde freshly dissolved from 50% stock in an artificial saline we strongly buffer to pH7.4 with 30mM HEPES buffer (and commensurately reduce NaCl to keep it isotonic). After two or three exchanges of this fixative at RT, and at least overnight fixation at RT (never at 4°C, because cold hurts membranes at this point), this primary fixative is washed away with at least two exchanges of 100mM cacodylate buffer, pH7.4 (since we have found that HEPES buffer, while gentle to cells, unfortunately suppresses the osmium-impregnation desired next).

The postfixation is conducted almost exactly as Karnovsky first proposed (Karnovsky, 1971), but because tissue cultures are so thin, and so permeable and gossimer, with the OsO4 reduced to 0.25% and the potassium ferrocyanide also reduced to 0.25%, and the duration of this postfixation reduced to 30min at RT. Thereafter, the cultures are washed again in cacodylate buffer. (In this and all steps, this cacodylate buffer contains 2mM CaCl2, which we have found assists greatly in the proper preservation of all cellular membranes.) Next, the culture is immersed in 1% tannic acid in 0.1M cacodylate buffer plus Ca++ (a modification of our own, of the original protocols for ’mordanting’ tissues with tannic acid (Tilney et al., 1973, Maupin and Pollard, 1983), which we realized is actually most useful for enhancing the uranyl acetate block-staining, which will follow). After 30min in tannic acid in cacodylate, they are again washed briefly in cacodylate buffer plus Ca++ and then dropped from pH 7.4 by a brief wash in 0.1M Na acetate buffer at pH5.2, in preparation for 30min of block-staining’ with 0.25% uranyl acetate in 50mM acetate buffer at pH5.2. Finally, after the UA is washed away with a brief rinse again in acetate buffer, they are progressively dehydrated through changes of 50% and 75% ethanol and then immersed in 100% ethanol (certified 200 proof).

At this point, we introduce the critically important step of transferring the glass-bottomed p35 culture-dishes into 1oz. polypropylene-plastic bottles, because our epoxy-embedding classically involves a propylene oxide (PO) step, in-between alcohol and plastic, but PO would dissolve the plastic culture dish *itself* and make a huge mess. Fortunately, we have found that overnight in 100% ethanol in the polypropylene-plastic bottles nicely softens the glue that attaches the glass to the bottoms of the p35 culture dishes, and allows us to separate them and remove and discard the culture-dish itself. Thereafter, the glass bottom, still in 100% ethanol, is rinsed for a few minutes in 100% propylene oxide and then embedded in epoxy in the usual way: a few hours in a mixture of 2/3rds epoxy and 1/3rd propylene oxide, then overnight at RT in 100% epoxy on a rotator, then transfer into freshly-prepared epoxy for the final vacuum-embedding, polymerizing this epoxy at 70°C for at least 24hr. We still use only the old English ’Araldite’ epoxy-formulation, finding it better than Epon for thin-sectioning and later grid-staining, and is still available from Polysciences, Ladd, or EMS.

For this whole embedding, including the polymerization, the glass is left in the polypropylene bottle which is inert to propylene oxide; but after hardening, the epoxy block can be popped easily out of the bottle, leaving the culture on the very bottom of a block of hardened epoxy, which can then be cut into small pieces appropriate for the chuck of an ultramicrotome. Just before this mounting in the microtome-chuck, the glass is finally removed from under the embedded culture by 5 sec contact with a brass block cooled to LN2 temperature (using the ’trick’ of the differential thermal-contraction of glass and plastic), or if that fails, as it sometimes does, by immersion of the block in full strength (47%) hydrofluoric acid, which completely dissolves away the glass in less than 10min, but does not penetrate the epoxy or degrade the cells in any way.

After thin-sectioning at 80nm (or at 40nm for ’crisper’ views of membranes, or at 150nm for eventual 3-D viewing), sections are picked up on Formvar/carbon coated 300 mesh ’wide view’ (e.g., narrow grid-bar) copper EM grids from Guilder, Ltd, and stained for 5 min with 1% lead citrate before viewing with a JEOL JEM1400 electron microscope, using its smallest objective aperture as well as a low 80KV voltage to achieve maximum contrast and clarity. Digital EM-images are captured at 6000 x 8000 pixel resolution with an AMT Biosprint 29 camera and manipulated or colorized when necessary (or made into 3-D anaglyphs’) with Adobe Photoshop, as described (Heuser, 2000a).

## ACKNOWLEDGMENTS

We wish to thank the following colleagues for providing us over many years with tissue cultures transfected with a whole variety of different reagents, for comparative purposes: Phyllis Hanson and Mark Diamond, then of Washington University School of Medicine in St. Louis; Azumi Yoshimura, Nobohiro Morone, Meneku Kengaku, and Silvia Pujals, then of Kyoto University, Japan; Jiin Felgner of UC Irvine; and Elena Mekhedov of NICHD, NIH.

## AUTHOR CONTRIBUTIONS

JP produced all the cultures and conducted all the transfection-experiments. TT did all of their preparation for EM, and all of their thin-sectioning. JH did all the electron microscopy, and wrote the whole manuscript. TB and PF monitored and advised at every step of the work, and extensively edited the manuscript.

---

**Supplementary Figure 1.**
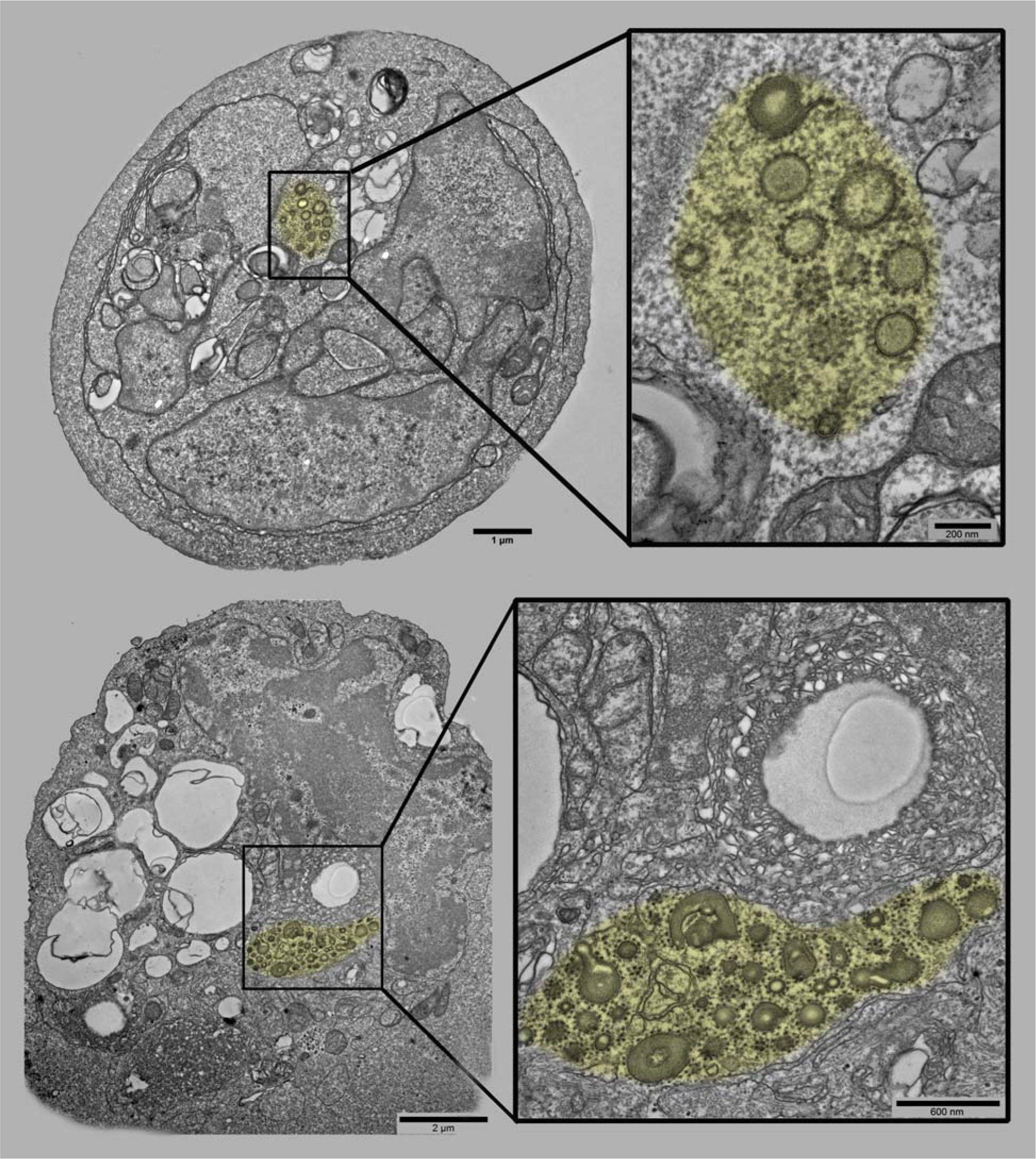
Two examples of the sorts of apoptotic cells invariably found in Lipofectamine-transfections, wherein the ribosome-studded clusters of lipofectamine-droplets remain present *throughout*, and remain in relatively well-defined and partitioned domains within the cytoplasm, domains that utterly exclude all other cytoplasmic organelles and thus can be considered to be separate *phases* in the cytoplasm (highlighted yellow). Scale bars indicate the relative magnification of the overviews, versus their respective enlarged insets.

**Supplementary Figure 2.**
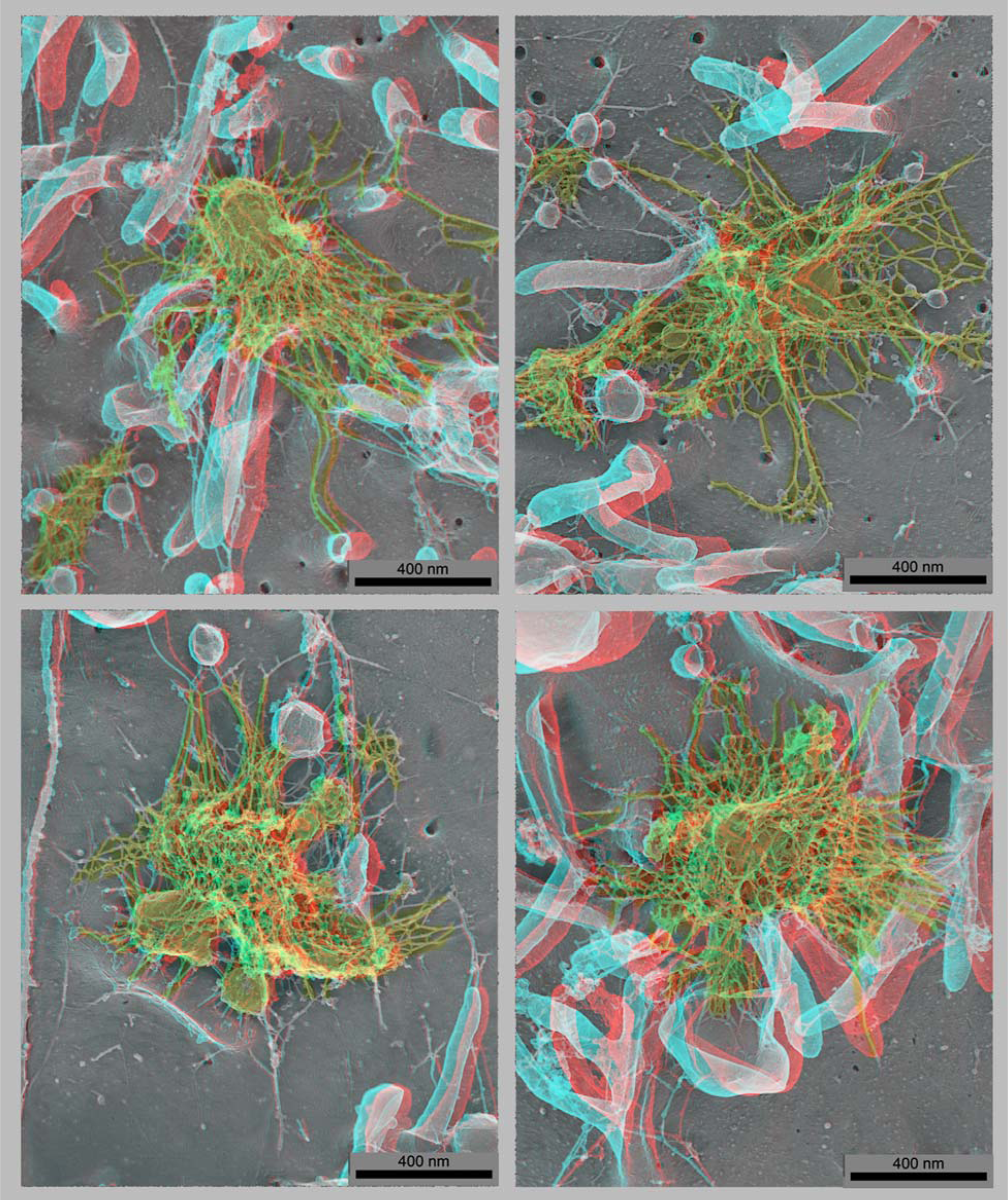
Four "anaglyph" 3-D views of the "deep-etch" EM-appearance of Lipofectamine+DNA complexes attached to the surfaces of cells. The lipoplexes as well as the thin ’hairs’ presumably representing their plasmid DNA are highlighted yellow. (Anaglyph-conversion of 3-D EM’s was applied as indicated in Heuser, 2000a.) Scale bars 400nm.

## REFERENCES (unformatted)

1. Afzelius, B.A. 1992. Section staining for electron microscopy using tannic acid as a mordant: a simple method for visualization of glycogen and collagen. Microsc Res Tech. 21:65–72.

2. Alberti, S., and A.A. Hyman. 2021. Biomolecular condensates at the nexus of cellular stress, protein aggregation disease and ageing. Nat Rev Mol Cell Biol. 22:196–213.

3. Asensio, C.S., D.W. Sirkis, J.W. Maas, Jr., K. Egami, T.L. To, F.M. Brodsky, X. Shu, Y. Cheng, and R.H. Edwards. 2013. Self-assembly of VPS41 promotes sorting required for biogenesis of the regulated secretory pathway. Dev Cell. 27:425–437.

4. Aytar, B.S., J.P. Muller, S. Golan, S. Hata, H. Takahashi, Y. Kondo, Y. Talmon, N.L. Abbott, and D.M. Lynn. 2012. Addition of ascorbic acid to the extracellular environment activates lipoplexes of a ferrocenyl lipid and promotes cell transfection. J Control Release. 157:249–259.

5. Baker, N.A., D. Sept, S. Joseph, M.J. Holst, and J.A. McCammon. 2001. Electrostatics of nanosystems: application to microtubules and the ribosome. Proc Natl Acad Sci U S A. 98:10037–10041.

6. Ballarin-Gonzalez, B., and K.A. Howard. 2012. Polycation-based nanoparticle delivery of RNAi therapeutics: adverse effects and solutions. Adv Drug Deliv Rev. 64:1717–1729.

7. Banani, S.F., H.O. Lee, A.A. Hyman, and M.K. Rosen. 2017. Biomolecular condensates: organizers of cellular biochemistry. Nat Rev Mol Cell Biol. 18:285–298.

8. Basha, G., T.I. Novobrantseva, N. Rosin, Y.Y. Tam, I.M. Hafez, M.K. Wong, T. Sugo, V.M. Ruda, J. Qin, B. Klebanov, M. Ciufolini, A. Akinc, Y.K. Tam, M.J. Hope, and P.R. Cullis. 2011. Influence of cationic lipid composition on gene silencing properties of lipid nanoparticle formulations of siRNA in antigen-presenting cells. Mol Ther. 19:2186–2200.

9. Battersby, B.J., R. Grimm, S. Huebner, and G. Cevc. 1998. Evidence for three-dimensional interlayer correlations in cationic lipid-DNA complexes as observed by cryo-electron microscopy. Biochim Biophys Acta. 1372:379–383.

10. Bauer, C., and P. Traub. 1995. Interaction of intermediate filaments with ribosomes in vitro. Eur J Cell Biol. 68:288–296.

11. Bauer, M., B.W. Kristensen, M. Meyer, T. Gasser, H.R. Widmer, J. Zimmer, and M. Ueffing. 2006. Toxic effects of lipid-mediated gene transfer in ventral mesencephalic explant cultures. Basic Clin Pharmacol Toxicol. 98:395–400.

12. Bawdon, R.E., A.M. Fiskin, and R.G. Garrison. 1984. Ultrastructural aspects of cytoplasmic ribosomes from Histoplasma capsulatum and Blastomyces dermatitidis as revealed by heavy metal staining. Mycopathologia. 86:155–163.

13. Beiras, A., T. Garcia-Caballero, R. Gallego, and E. Roson. 1987. Staining of neuroendocrine Merkel cells of human epidermis using the uranaffin reaction. J Invest Dermatol. 89:366–368.

14. Bernhard, W. 1969. A new staining procedure for electron microscopical cytology. J Ultrastruct Res. 27:250–265.

15. Biazik, J., H. Vihinen, E. Jokitalo, and E.L. Eskelinen. 2017. Ultrastructural Characterization of Phagophores Using Electron Tomography on Cryoimmobilized and Freeze Substituted Samples. Methods Enzymol. 587:331–349.

16. Biazik, J., P. Yla-Anttila, H. Vihinen, E. Jokitalo, and E.L. Eskelinen. 2015. Ultrastructural relationship of the phagophore with surrounding organelles. Autophagy. 11:439–451.

17. Blad, D., L. Winqvist, and G. Dallner. 1977. Electrophoretic mobility of microsomes from rat liver. J Cell Sci. 23:285–297.

18. Blad-Holmberg, D. 1979. Study of electrophoretic mobility of cellular membranes isolated from rat liver. Biochim Biophys Acta. 553:25–39.

19. Boassa, D., M.L. Berlanga, M.A. Yang, M. Terada, J. Hu, E.A. Bushong, M. Hwang, E. Masliah, J.M. George, and M.H. Ellisman. 2013. Mapping the subcellular distribution of alpha-synuclein in neurons using genetically encoded probes for correlated light and electron microscopy: implications for Parkinson’s disease pathogenesis. J Neurosci. 33:2605–2615.

20. Borgese, N., N. Iacomino, S.F. Colombo, and F. Navone. 2021. The Link between VAPB Loss of Function and Amyotrophic Lateral Sclerosis. Cells. 10.

21. Brangwynne, C.P., C.R. Eckmann, D.S. Courson, A. Rybarska, C. Hoege, J. Gharakhani, F. Julicher, and A.A. Hyman. 2009. Germline P granules are liquid droplets that localize by controlled dissolution/condensation. Science. 324:1729–1732.

22. Brodie, D.A., P. Huie, M. Locke, and F.P. Ottensmeyer. 1982. The correlation between bismuth and uranyl staining and phosphorus content of intracellular structures as determined by electron spectroscopic imaging. Tissue Cell. 14:621–627.

23. Brunner, S., T. Sauer, S. Carotta, M. Cotten, M. Saltik, and E. Wagner. 2000. Cell cycle dependence of gene transfer by lipoplex, polyplex and recombinant adenovirus. Gene Ther. 7:401–407.

24. Caracciolo, G., R. Caminiti, M.A. Digman, E. Gratton, and S. Sanchez. 2009. Efficient escape from endosomes determines the superior efficiency of multicomponent lipoplexes. J Phys Chem B. 113:4995–4997.

25. Carter SD, Hampton CM, Langlois R, Melero R, Farino ZJ, Calderon MJ, Li W, Wallace CT, Tran NH, Grassucci RA, Siegmund SE, Pemberton J, Morgenstern TJ, Eisenman L, Aguilar JI, Greenberg NL, Levy ES, Yi E, Mitchell WG, Rice WJ, Wigge C, Pilli J, George EW, Aslanoglou D, Courel M, Freyberg RJ, Javitch JA, Wills ZP, Area-Gomez E, Shiva S, Bartolini F, Volchuk A, Murray SA, Aridor M, Fish KN, Walter P, Balla T, Fass D, Wolf SG, Watkins SC, Carazo JM, Jensen GJ, Frank J, Freyberg Z. 2020. Ribosome-associated vesicles: A dynamic subcompartment of the endoplasmic reticulum in secretory cells. Sci Adv. 6: eaay9572 Chernousova, S., and M. Epple. 2017. Live-cell imaging to compare the transfection and gene silencing efficiency of calcium phosphate nanoparticles and a liposomal transfection agent. Gene Ther. 24:282–289.

26. Colin, M., M. Maurice, G. Trugnan, M. Kornprobst, R.P. Harbottle, A. Knight, R.G. Cooper, A.D. Miller, J. Capeau, C. Coutelle, and M.C. Brahimi-Horn. 2000. Cell delivery, intracellular trafficking and expression of an integrin-mediated gene transfer vector in tracheal epithelial cells. Gene Ther. 7:139–152.

27. Cordes, V.C., S. Reidenbach, and W.W. Franke. 1996. Cytoplasmic annulate lamellae in cultured cells: composition, distribution, and mitotic behavior. Cell Tissue Res. 284:177–191.

28. Dalby, B., S. Cates, A. Harris, E.C. Ohki, M.L. Tilkins, P.J. Price, and V.C. Ciccarone. 2004. Advanced transfection with Lipofectamine 2000 reagent: primary neurons, siRNA, and high-throughput applications. Methods. 33:95–103.

29. Dallner, G., P. Siekevitz, and G.E. Palade. 1966. Biogenesis of endoplasmic reticulum membranes. I. Structural and chemical differentiation in developing rat hepatocyte. J Cell Biol. 30:73–96.

30. Damodaran, A.P., T. Courtheoux, E. Watrin, and C. Prigent. 2020. Alteration of SC35 localization by transfection reagents. Biochim Biophys Acta Mol Cell Res. 1867:118650.

31. Dokka, S., D. Toledo, X. Shi, V. Castranova, and Y. Rojanasakul. 2000. Oxygen radical-mediated pulmonary toxicity induced by some cationic liposomes. Pharm Res. 17:521–525.

32. Donkuru, M., I. Badea, S. Wettig, R. Verrall, M. Elsabahy, and M. Foldvari. 2010. Advancing nonviral gene delivery: lipid- and surfactant-based nanoparticle design strategies. Nanomedicine (Lond). 5:1103–1127.

33. Dunlap, D.D., A. Maggi, M.R. Soria, and L. Monaco. 1997. Nanoscopic structure of DNA condensed for gene delivery. Nucleic Acids Res. 25:3095–3101.

34. Duzgunes, N., and P.L. Felgner. 1993. Intracellular delivery of nucleic acids and transcription factors by cationic liposomes. Methods Enzymol. 221:303–306.

35. Duzgunes, N., J.A. Goldstein, D.S. Friend, and P.L. Felgner. 1989. Fusion of liposomes containing a novel cationic lipid, N-[2,3-(dioleyloxy)propyl]-N,N,N-trimethylammonium: induction by multivalent anions and asymmetric fusion with acidic phospholipid vesicles. Biochemistry. 28:9179–9184.

36. Dvorak, A.M., and E.S. Morgan. 2001. Ribosomes and secretory granules in human mast cells: close associations demonstrated by staining with a chelating agent. Immunol Rev. 179:94–101.

37. Erenpreisa, J., A. Ivanov, M. Cragg, G. Selivanova, and T. Illidge. 2002. Nuclear envelope-limited chromatin sheets are part of mitotic death. Histochem Cell Biol. 117:243–255.

38. Ewert, K., H.M. Evans, A. Ahmad, N.L. Slack, A.J. Lin, A. Martin-Herranz, and C.R. Safinya. 2005. Lipoplex structures and their distinct cellular pathways. Adv Genet. 53:119–155.

39. Ewert, K.K., P. Scodeller, L. Simon-Gracia, V.M. Steffes, E.A. Wonder, T. Teesalu, and C.R. Safinya. 2021. Cationic Liposomes as Vectors for Nucleic Acid and Hydrophobic Drug Therapeutics. Pharmaceutics. 13.

40. Eymieux, S., Y. Rouille, O. Terrier, K. Seron, E. Blanchard, M. Rosa-Calatrava, J. Dubuisson, S. Belouzard, and P. Roingeard. 2021. Ultrastructural modifications induced by SARS-CoV-2 in Vero cells: a kinetic analysis of viral factory formation, viral particle morphogenesis and virion release. Cell Mol Life Sci. 78:3565–3576.

41. Farquhar, M.G., and G.E. Palade. 1965. Cell junctions in amphibian skin. J Cell Biol. 26:263–291.

42. Fasana, E., M. Fossati, A. Ruggiano, S. Brambillasca, C.C. Hoogenraad, F. Navone, M. Francolini, and N. Borgese. 2010. A VAPB mutant linked to amyotrophic lateral sclerosis generates a novel form of organized smooth endoplasmic reticulum. FASEB J. 24:1419–1430.

43. Felgner, J.H., R. Kumar, C.N. Sridhar, C.J. Wheeler, Y.J. Tsai, R. Border, P. Ramsey, M. Martin, and P.L. Felgner. 1994. Enhanced gene delivery and mechanism studies with a novel series of cationic lipid formulations. J Biol Chem. 269:2550–2561.

44. Felgner, P.L. 1997. Nonviral strategies for gene therapy. Sci Am. 276:102–106.

45. Felgner, P.L. 1999. Prospects for synthetic self-assembling systems in gene delivery. J Gene Med. 1:290–292.

46. Felgner, P.L., T.R. Gadek, M. Holm, R. Roman, H.W. Chan, M. Wenz, J.P. Northrop, G.M. Ringold, and M. Danielsen. 1987. Lipofection: a highly efficient, lipid-mediated DNA-transfection procedure. Proc Natl Acad Sci U S A. 84:7413–7417.

47. Fengsrud, M., N. Roos, T. Berg, W. Liou, J.W. Slot, and P.O. Seglen. 1995. Ultrastructural and immunocytochemical characterization of autophagic vacuoles in isolated hepatocytes: effects of vinblastine and asparagine on vacuole distributions. Exp Cell Res. 221:504–519.

48. Fenske, D.B., and P.R. Cullis. 2005. Entrapment of small molecules and nucleic acid-based drugs in liposomes. Methods Enzymol. 391:7–40.

49. Filion, M.C., and N.C. Phillips. 1997. Toxicity and immunomodulatory activity of liposomal vectors formulated with cationic lipids toward immune effector cells. Biochim Biophys Acta. 1329:345–356.

50. Friend, D.S., D. Papahadjopoulos, and R.J. Debs. 1996. Endocytosis and intracellular processing accompanying transfection mediated by cationic liposomes. Biochim Biophys Acta. 1278:41–50.

51. Frost, B., R.L. Jacks, and M.I. Diamond. 2009. Propagation of tau misfolding from the outside to the inside of a cell. J Biol Chem. 284:12845–12852.

52. Goryaynov, A., and W. Yang. 2014. Role of molecular charge in nucleocytoplasmic transport. PLoS One. 9:e88792.

53. Gudmundsson, S.R., K.A. Kallio, H. Vihinen, E. Jokitalo, N. Ktistakis, and E.L. Eskelinen. 2022. Morphology of Phagophore Precursors by Correlative Light-Electron Microscopy. Cells. 11.

54. Gustafsson, J., G. Arvidson, G. Karlsson, and M. Almgren. 1995. Complexes between cationic liposomes and DNA visualized by cryo-TEM. Biochim Biophys Acta. 1235:305–312.

55. Haggag, A., and J. Gilloteaux. 2005. Unusual circular annulate lamellae in hepatocytes of Torpedo marmorata. Histol Histopathol. 20:785–789.

56. Hanson, H.H., S. Kang, M. Fernandez-Monreal, T. Oung, M. Yildirim, R. Lee, K. Suyama, R.B. Hazan, and G.R. Phillips. 2010. LC3-dependent intracellular membrane tubules induced by gamma-protocadherins A3 and B2: a role for intraluminal interactions. J Biol Chem. 285:20982–20992.

57. Hawley-Nelson, P., and V. Ciccarone. 2001. Transfection of cultured eukaryotic cells using cationic lipid reagents. Curr Protoc Neurosci. Appendix 1:Appendix 1F.

58. Hawley-Nelson, P., and V. Ciccarone. 2003. Transfection of cultured eukaryotic cells using cationic lipid reagents. Curr Protoc Cell Biol. Chapter 20:Unit 20 26.

59. Hawley-Nelson, P., V. Ciccarone, and M.L. Moore. 2008. Transfection of cultured eukaryotic cells using cationic lipid reagents. Curr Protoc Mol Biol. Chapter 9:Unit 9 4.

60. Hayashi-Nishino, M., N. Fujita, T. Noda, A. Yamaguchi, T. Yoshimori, and A. Yamamoto. 2009. A subdomain of the endoplasmic reticulum forms a cradle for autophagosome formation. Nat Cell Biol. 11:1433–1437.

61. Heuser, J. 1981. Preparing biological samples for stereomicroscopy by the quick-freeze, deep-etch, rotary-replication technique. Methods Cell Biol. 22:97–122.

62. Heuser, J. 1989a. Protocol for 3-D visualization of molecules on mica via the quick-freeze, deep-etch technique. J Electron Microsc Tech. 13:244–263.

63. Heuser, J. 2000a. The production of ’cell cortices’ for light and electron microscopy. Traffic. 1:545–552.

64. Heuser, J.E. 1983. Procedure for freeze-drying molecules adsorbed to mica flakes. J Mol Biol. 169:155–195.

65. Heuser, J.E. 1989b. Development of the quick-freeze, deep-etch, rotary-replication technique of sample preparation for 3-D electron microscopy. Prog Clin Biol Res. 295:71–83.

66. Heuser, J.E. 2000b. Membrane traffic in anaglyph stereo. Traffic. 1:35–37.

67. Heuser, J.E. 2011. The origins and evolution of freeze-etch electron microscopy. J Electron Microsc (Tokyo). 60 Suppl 1:S3–29.

68. Heuser, J.E. 2022. The Structural Basis of Long-Term Potentiation in Hippocampal Synapses, Revealed by Electron Microscopy Imaging of Lanthanum-Induced Synaptic Vesicle Recycling. Front Cell Neurosci. 16:920360.

69. Heuser, J.E., T.S. Reese, M.J. Dennis, Y. Jan, L. Jan, and L. Evans. 1979. Synaptic vesicle exocytosis captured by quick freezing and correlated with quantal transmitter release. J Cell Biol. 81:275–300.

70. Heuser, J.E., and S.R. Salpeter. 1979. Organization of acetylcholine receptors in quick-frozen, deep-etched, and rotary-replicated Torpedo postsynaptic membrane. J Cell Biol. 82:150–173.

71. Heuser, J.E., and T.I. Tenkova. 2020. Introducing a mammalian nerve-muscle preparation ideal for physiology and microscopy, the transverse auricular muscle in the ear of the mouse. Neuroscience. 439:80–105.

72. Hoekstra, D., J. Rejman, L. Wasungu, F. Shi, and I. Zuhorn. 2007. Gene delivery by cationic lipids: in and out of an endosome. Biochem Soc Trans. 35:68–71.

73. Holmes, B.B., S.L. DeVos, N. Kfoury, M. Li, R. Jacks, K. Yanamandra, M.O. Ouidja, F.M. Brodsky, J. Marasa, D.P. Bagchi, P.T. Kotzbauer, T.M. Miller, D. Papy-Garcia, and M.I. Diamond. 2013. Heparan sulfate proteoglycans mediate internalization and propagation of specific proteopathic seeds. Proc Natl Acad Sci U S A. 110:E3138–3147.

74. Huber, S., A. Bar, S. Epp, J. Schmuckli-Maurer, N. Eberhard, B.M. Humbel, A. Hemphill, and K. Woods. 2020. Recruitment of Host Nuclear Pore Components to the Vicinity of Theileria Schizonts. mSphere. 5.

75. Huebner, S., B.J. Battersby, R. Grimm, and G. Cevc. 1999. Lipid-DNA complex formation: reorganization and rupture of lipid vesicles in the presence of DNA as observed by cryoelectron microscopy. Biophys J. 76:3158–3166.

76. Huxley, H.E., and G. Zubay. 1961. Preferential staining of nucleic acid-containing structures for electron microscopy. J Biophys Biochem Cytol. 11:273–296.

77. Hyman, A.A., and C.P. Brangwynne. 2011. Beyond stereospecificity: liquids and mesoscale organization of cytoplasm. Dev Cell. 21:14–16.

78. Hyman, A.A., C.A. Weber, and F. Julicher. 2014. Liquid-liquid phase separation in biology. Annu Rev Cell Dev Biol. 30:39–58.

79. Imreh, G., and E. Hallberg. 2000. An integral membrane protein from the nuclear pore complex is also present in the annulate lamellae: implications for annulate lamella formation. Exp Cell Res. 259:180–190.

80. Kanekura, K., I. Nishimoto, S. Aiso, and M. Matsuoka. 2006. Characterization of amyotrophic lateral sclerosis-linked P56S mutation of vesicle-associated membrane protein-associated protein B (VAPB/ALS8). J Biol Chem. 281:30223–30233.

81. Kanekura, K., H. Suzuki, S. Aiso, and M. Matsuoka. 2009. ER stress and unfolded protein response in amyotrophic lateral sclerosis. Mol Neurobiol. 39:81–89.

82. Karnovsky, M.J. 1961. Simple methods for "staining with lead" at high pH in electron microscopy. J Biophys Biochem Cytol. 11:729–732.

83. Karnovsky, M.J. 1971. Use of ferrocyanide-reduced osmium tetroxide in electron microscopy. In Abstracts of the American Society of Cell Biology. New Orleans. 146.

84. Kawabuchi, M., M. Osame, Y. Aika, and T. Kanaseki. 1989. Annulate lamellae-soleplate nuclei associations in skeletal muscle fibers of rats during chronic high-dose exposure to neostigmine. Anat Rec. 225:1–10.

85. Kessel, R.G. 1983a. Fibrogranular bodies, annulate lamellae, and polyribosomes in the dragonfly oocyte. J Morphol. 176:171–180.

86. Kessel, R.G. 1983b. The structure and function of annulate lamellae: porous cytoplasmic and intranuclear membranes. Int Rev Cytol. 82:181–303.

87. Kfoury, N., B.B. Holmes, H. Jiang, D.M. Holtzman, and M.I. Diamond. 2012. Trans-cellular propagation of Tau aggregation by fibrillar species. J Biol Chem. 287:19440–19451.

88. Kislev, N., H. Swift, and L. Bogorad. 1965. Nucleic Acids of Chloroplasts and Mitochondria in Swiss Chard. J Cell Biol. 25:327–344.

89. Knight, A.M., P.H. Culviner, N. Kurt-Yilmaz, T. Zou, S.B. Ozkan, and S. Cavagnero. 2013. Electrostatic effect of the ribosomal surface on nascent polypeptide dynamics. ACS Chem Biol. 8:1195–1204.

90. Kolay, S., and M.I. Diamond. 2020. Alzheimer’s disease risk modifier genes do not affect tau aggregate uptake, seeding or maintenance in cell models. FEBS Open Bio. 10:1912–1920.

91. Koltover, I., T. Salditt, J.O. Radler, and C.R. Safinya. 1998. An inverted hexagonal phase of cationic liposome-DNA complexes related to DNA release and delivery. Science. 281:78–81.

92. Konopka, K., G.S. Harrison, P.L. Felgner, and N. Duzgunes. 1997. Cationic liposome-mediated expression of HIV-regulated luciferase and diphtheria toxin a genes in HeLa cells infected with or expressing HIV. Biochim Biophys Acta. 1356:185–197.

93. Konopka, K., E. Pretzer, P.L. Felgner, and N. Duzgunes. 1996. Human immunodeficiency virus type-1 (HIV-1) infection increases the sensitivity of macrophages and THP-1 cells to cytotoxicity by cationic liposomes. Biochim Biophys Acta. 1312:186–196.

94. Korn, A.P., P. Spitnik-Elson, and D. Elson. 1985. Identification of the collar-like structure of the 30S ribosomal subunit from E. coli by dark field electron microscopy. Eur J Cell Biol. 39:56–61.

95. Kovacs, A.L., A. Reith, and P.O. Seglen. 1982. Accumulation of autophagosomes after inhibition of hepatocytic protein degradation by vinblastine, leupeptin or a lysosomotropic amine. Exp Cell Res. 137:191–201.

96. Kuijpers, M., V. van Dis, E.D. Haasdijk, M. Harterink, K. Vocking, J.A. Post, W. Scheper, C.C. Hoogenraad, and D. Jaarsma. 2013. Amyotrophic lateral sclerosis (ALS)-associated VAPB- P56S inclusions represent an ER quality control compartment. Acta Neuropathol Commun. 1:24.

97. Kulkarni, J.A., M.M. Darjuan, J.E. Mercer, S. Chen, R. van der Meel, J.L. Thewalt, Y.Y.C. Tam, and P.R. Cullis. 2018. On the Formation and Morphology of Lipid Nanoparticles Containing Ionizable Cationic Lipids and siRNA. ACS Nano. 12:4787–4795.

98. Kulkarni, J.A., S.B. Thomson, J. Zaifman, J. Leung, P.K. Wagner, A. Hill, Y.Y.C. Tam, P.R. Cullis, T.L. Petkau, and B.R. Leavitt. 2020. Spontaneous, solvent-free entrapment of siRNA within lipid nanoparticles. Nanoscale. 12:23959–23966.

99. Kulkarni, J.A., D. Witzigmann, S. Chen, P.R. Cullis, and R. van der Meel. 2019a. Lipid Nanoparticle Technology for Clinical Translation of siRNA Therapeutics. Acc Chem Res. 52:2435–2444.

100. Kulkarni, J.A., D. Witzigmann, J. Leung, R. van der Meel, J. Zaifman, M.M. Darjuan, H.M. Grisch-Chan, B. Thony, Y.Y.C. Tam, and P.R. Cullis. 2019b. Fusion-dependent formation of lipid nanoparticles containing macromolecular payloads. Nanoscale. 11:9023–9031.

101. Kuntsche, J., J.C. Horst, and H. Bunjes. 2011. Cryogenic transmission electron microscopy (cryo-TEM) for studying the morphology of colloidal drug delivery systems. Int J Pharm. 417:120–137.

102. Lazebnik, M., R.K. Keswani, and D.W. Pack. 2016. Endocytic Transport of Polyplex and Lipoplex siRNA Vectors in HeLa Cells. Pharm Res. 33:2999–3011.

103. Le Bihan, O., R. Chevre, S. Mornet, B. Garnier, B. Pitard, and O. Lambert. 2011. Probing the in vitro mechanism of action of cationic lipid/DNA lipoplexes at a nanometric scale. Nucleic Acids Res. 39:1595–1609.

104. Leal, C., N.F. Bouxsein, K.K. Ewert, and C.R. Safinya. 2010. Highly efficient gene silencing activity of siRNA embedded in a nanostructured gyroid cubic lipid matrix. J Am Chem Soc. 132:16841–16847.

105. Leung, A.K., Y.Y. Tam, S. Chen, I.M. Hafez, and P.R. Cullis. 2015. Microfluidic Mixing: A General Method for Encapsulating Macromolecules in Lipid Nanoparticle Systems. J Phys Chem B. 119:8698–8706.

106. Li, Z., C. Zhang, Z. Wang, J. Shen, P. Xiang, X. Chen, J. Nan, and Y. Lin. 2019. Lipofectamine 2000/siRNA complexes cause endoplasmic reticulum unfolded protein response in human endothelial cells. J Cell Physiol. 234:21166–21181.

107. Lin, A.J., N.L. Slack, A. Ahmad, C.X. George, C.E. Samuel, and C.R. Safinya. 2003. Three-dimensional imaging of lipid gene-carriers: membrane charge density controls universal transfection behavior in lamellar cationic liposome-DNA complexes. Biophys J. 84:3307–3316.

108. Lin, D.H., and A. Hoelz. 2019. The Structure of the Nuclear Pore Complex (An Update). Annu Rev Biochem. 88:725–783.

109. Lin, Q., J. Chen, Z. Zhang, and G. Zheng. 2014. Lipid-based nanoparticles in the systemic delivery of siRNA. Nanomedicine (Lond). 9:105–120.

110. Liu, Y., L. Wenning, M. Lynch, and T.M. Reineke. 2004. New poly(d-glucaramidoamine)s induce DNA nanoparticle formation and efficient gene delivery into mammalian cells. J Am Chem Soc. 126:7422–7423.

111. Lozhkin, A., A.E. Vendrov, H. Pan, S.A. Wickline, N.R. Madamanchi, and M.S. Runge. 2017. NADPH oxidase 4 regulates vascular inflammation in aging and atherosclerosis. J Mol Cell Cardiol. 102:10–21.

112. Luk, K.C., C. Song, P. O’Brien, A. Stieber, J.R. Branch, K.R. Brunden, J.Q. Trojanowski, and V.M. Lee. 2009. Exogenous alpha-synuclein fibrils seed the formation of Lewy body-like intracellular inclusions in cultured cells. Proc Natl Acad Sci U S A. 106:20051–20056.

113. Lv, H., S. Zhang, B. Wang, S. Cui, and J. Yan. 2006. Toxicity of cationic lipids and cationic polymers in gene delivery. J Control Release. 114:100–109.

114. Magalhaes, S., S. Duarte, G.A. Monteiro, and F. Fernandes. 2014. Quantitative evaluation of DNA dissociation from liposome carriers and DNA escape from endosomes during lipid-mediated gene delivery. Hum Gene Ther Methods. 25:303–313.

115. Mahamid, J., S. Pfeffer, M. Schaffer, E. Villa, R. Danev, L.K. Cuellar, F. Forster, A.A. Hyman, J.M. Plitzko, and W. Baumeister. 2016. Visualizing the molecular sociology at the HeLa cell nuclear periphery. Science. 351:969–972.

116. Majzoub, R.N., C.L. Chan, K.K. Ewert, B.F. Silva, K.S. Liang, and C.R. Safinya. 2015. Fluorescence microscopy colocalization of lipid-nucleic acid nanoparticles with wildtype and mutant Rab5-GFP: A platform for investigating early endosomal events. Biochim Biophys Acta. 1848:1308–1318.

117. Majzoub, R.N., K.K. Ewert, and C.R. Safinya. 2016a. Cationic liposome-nucleic acid nanoparticle assemblies with applications in gene delivery and gene silencing. Philos Trans A Math Phys Eng Sci. 374.

118. Majzoub, R.N., K.K. Ewert, and C.R. Safinya. 2016b. Quantitative Intracellular Localization of Cationic Lipid-Nucleic Acid Nanoparticles with Fluorescence Microscopy. Methods Mol Biol. 1445:77–108.

119. Malatesta, M. 2021. Transmission Electron Microscopy as a Powerful Tool to Investigate the Interaction of Nanoparticles with Subcellular Structures. Int J Mol Sci. 22.

120. Malone, R.W., P.L. Felgner, and I.M. Verma. 1989. Cationic liposome-mediated RNA transfection. Proc Natl Acad Sci U S A. 86:6077–6081.

121. Martin, T.M., B.J. Wysocki, J.P. Beyersdorf, T.A. Wysocki, and A.K. Pannier. 2014. Integrating mitosis, toxicity, and transgene expression in a telecommunications packet-switched network model of lipoplex-mediated gene delivery. Biotechnol Bioeng. 111:1659–1671.

122. Maupin P, & TD Pollard (1983) Improved preservation and staining of cytoplasmic structures by tannic acid-glutaraldehyde-saponin fixation. J Cell Biol. 96: 51–62.

123. Maurer, N., A. Mori, L. Palmer, M.A. Monck, K.W. Mok, B. Mui, Q.F. Akhong, and P.R. Cullis. 1999. Lipid-based systems for the intracellular delivery of genetic drugs. Mol Membr Biol. 16:129–140.

124. Midoux, P., C. Pichon, J.J. Yaouanc, and P.A. Jaffres. 2009. Chemical vectors for gene delivery: a current review on polymers, peptides and lipids containing histidine or imidazole as nucleic acids carriers. Br J Pharmacol. 157:166–178.

125. Mo, R.H., J.L. Zaro, J.H. Ou, and W.C. Shen. 2012. Effects of Lipofectamine 2000/siRNA complexes on autophagy in hepatoma cells. Mol Biotechnol. 51:1–8.

126. Morgenstern, E. 1977. [Cytochemical studies of ultrathin frozen sections of blood platelets]. Acta Histochem Suppl. Suppl 18:259–264.

127. Napoli, E., S. Liu, I. Marsilio, K. Zarbalis, and C. Giulivi. 2017. Lipid-based DNA/siRNA transfection agents disrupt neuronal bioenergetics and mitophagy. Biochem J. 474:3887–3902.

128. Navone, F., P. Genevini, and N. Borgese. 2015. Autophagy and Neurodegeneration: Insights from a Cultured Cell Model of ALS. Cells. 4:354–386.

129. Nekooki-Machida, Y., M. Kurosawa, N. Nukina, K. Ito, T. Oda, and M. Tanaka. 2009. Distinct conformations of in vitro and in vivo amyloids of huntingtin-exon1 show different cytotoxicity. Proc Natl Acad Sci U S A. 106:9679–9684.

130. Nishimura, A.L., M. Mitne-Neto, H.C. Silva, A. Richieri-Costa, S. Middleton, D. Cascio, F. Kok, J.R. Oliveira, T. Gillingwater, J. Webb, P. Skehel, and M. Zatz. 2004. A mutation in the vesicle-trafficking protein VAPB causes late-onset spinal muscular atrophy and amyotrophic lateral sclerosis. Am J Hum Genet. 75:822–831.

131. Niso-Santano, M., J.M. Bravo-San Pedro, R. Gomez-Sanchez, V. Climent, G. Soler, J.M. Fuentes, and R.A. Gonzalez-Polo. 2011. ASK1 overexpression accelerates paraquat-induced autophagy via endoplasmic reticulum stress. Toxicol Sci. 119:156–168.

132. Ohki, E.C., M.L. Tilkins, V.C. Ciccarone, and P.J. Price. 2001. Improving the transfection efficiency of post-mitotic neurons. J Neurosci Methods. 112:95–99.

133. Papiani, G., A. Ruggiano, M. Fossati, A. Raimondi, G. Bertoni, M. Francolini, R. Benfante, F. Navone, and N. Borgese. 2012. Restructured endoplasmic reticulum generated by mutant amyotrophic lateral sclerosis-linked VAPB is cleared by the proteasome. J Cell Sci. 125:3601–3611.

134. Park, S., C. Zuber, and J. Roth. 2020. Selective autophagy of cytosolic protein aggregates involves ribosome-free rough endoplasmic reticulum. Histochem Cell Biol. 153:89–99.

135. Payne, C.M., R.B. Nagle, V.F. Borduin, and A. Kim. 1983. An ultrastructural evaluation of the cell organelle specificity of the uranaffin reaction in two human endocrine neoplasms. J Submicrosc Cytol. 15:833–841.

136. Piefke, J., T. Arad, H.S. Gewitz, A. Yonath, and H.G. Wittmann. 1986. The growth of ordered two-dimensional sheets of 70 S ribosomes from Bacillus stearothermophilus. FEBS Lett. 209:104–106.

137. Plaza-Ga, I., V. Manzaneda-Gonzalez, M. Kisovec, V. Almendro-Vedia, M. Munoz-Ubeda, G. Anderluh, A. Guerrero-Martinez, P. Natale, and I. Lopez Montero. 2019. pH-triggered endosomal escape of pore-forming Listeriolysin O toxin-coated gold nanoparticles. J Nanobiotechnology. 17:108.

138. Radler, J.O., I. Koltover, T. Salditt, and C.R. Safinya. 1997. Structure of DNA-cationic liposome complexes: DNA intercalation in multilamellar membranes in distinct interhelical packing regimes. Science. 275:810–814.

139. Raghunayakula, S., D. Subramonian, M. Dasso, R. Kumar, and X.D. Zhang. 2015. Molecular Characterization and Functional Analysis of Annulate Lamellae Pore Complexes in Nuclear Transport in Mammalian Cells. PLoS One. 10:e0144508.

140. Ramezanpour, M., M.L. Schmidt, I. Bodnariuc, J.A. Kulkarni, S.S.W. Leung, P.R. Cullis, J.L. Thewalt, and D.P. Tieleman. 2019. Ionizable amino lipid interactions with POPC: implications for lipid nanoparticle function. Nanoscale. 11:14141–14146.

141. Reifarth, M., S. Hoeppener, and U.S. Schubert. 2018. Uptake and Intracellular Fate of Engineered Nanoparticles in Mammalian Cells: Capabilities and Limitations of Transmission Electron Microscopy-Polymer-Based Nanoparticles. Adv Mater. 30.

142. Reynolds, E.S. 1963. The use of lead citrate at high pH as an electron-opaque stain in electron microscopy. J Cell Biol. 17:208–212.

143. Safinya, C.R., K.K. Ewert, R.N. Majzoub, and C. Leal. 2014. Cationic liposome-nucleic acid complexes for gene delivery and gene silencing. New J Chem. 38:5164–5172.

144. Sanders, D.W., S.K. Kaufman, S.L. DeVos, A.M. Sharma, H. Mirbaha, A. Li, S.J. Barker, A.C. Foley, J.R. Thorpe, L.C. Serpell, T.M. Miller, L.T. Grinberg, W.W. Seeley, and M.I. Diamond. 2014. Distinct tau prion strains propagate in cells and mice and define different tauopathies. Neuron. 82:1271–1288.

145. Sawaguchi, A., S. Ide, Y. Goto, J. Kawano, T. Oinuma, and T. Suganuma. 2001. A simple contrast enhancement by potassium permanganate oxidation for Lowicryl K4M ultrathin sections prepared by high pressure freezing/freeze substitution. J Microsc. 201:77–83.

146. Schavemaker, P.E., W.M. Smigiel, and B. Poolman. 2017. Ribosome surface properties may impose limits on the nature of the cytoplasmic proteome. Elife. 6.

147. Schiavone, N., L. Papucci, P. Luciani, A. Lapucci, M. Donnini, and S. Capaccioli. 2000. Induction of apoptosis and mitosis inhibition by degraded DNA lipotransfection mimicking genotoxic drug effects. Biochem Biophys Res Commun. 270:406–414.

148. Shu, X., V. Lev-Ram, T.J. Deerinck, Y. Qi, E.B. Ramko, M.W. Davidson, Y. Jin, M.H. Ellisman, and R.Y. Tsien. 2011. A genetically encoded tag for correlated light and electron microscopy of intact cells, tissues, and organisms. PLoS Biol. 9:e1001041.

149. Silva, J.P., I.M. Oliveira, A.C. Oliveira, M. Lucio, A.C. Gomes, P.J. Coutinho, and M.E. Oliveira. 2014. Structural dynamics and physicochemical properties of pDNA/DODAB:MO lipoplexes: effect of pH and anionic lipids in inverted non-lamellar phases versus lamellar phases. Biochim Biophys Acta. 1838:2555–2567.

150. Singh, R.S., C. Goncalves, P. Sandrin, C. Pichon, P. Midoux, and A. Chaudhuri. 2004. On the gene delivery efficacies of pH-sensitive cationic lipids via endosomal protonation: a chemical biology investigation. Chem Biol. 11:713–723.

151. Soloff, B.L. 1973. Buffered potassium permanganate-uranyl acetate-lead citrate staining seguence for ultrathin sections. Stain Technol. 48:159–165.

152. Stephan, D.J., Z.Y. Yang, H. San, R.D. Simari, C.J. Wheeler, P.L. Felgner, D. Gordon, G.J. Nabel, and E.G. Nabel. 1996. A new cationic liposome DNA complex enhances the efficiency of arterial gene transfer in vivo. Hum Gene Ther. 7:1803–1812.

153. Sternberg, B., K. Hong, W. Zheng, and D. Papahadjopoulos. 1998. Ultrastructural characterization of cationic liposome-DNA complexes showing enhanced stability in serum and high transfection activity in vivo. Biochim Biophys Acta. 1375:23–35.

154. Sternberg, B., F.L. Sorgi, and L. Huang. 1994. New structures in complex formation between DNA and cationic liposomes visualized by freeze-fracture electron microscopy. FEBS Lett. 356:361–366.

155. Terada, T., J.A. Kulkarni, A. Huynh, S. Chen, R. van der Meel, Y.Y.C. Tam, and P.R. Cullis. 2021a. Characterization of Lipid Nanoparticles Containing Ionizable Cationic Lipids Using Design-of-Experiments Approach. Langmuir. 37:1120–1128.

156. Terada, T., J.A. Kulkarni, A. Huynh, Y.Y.C. Tam, and P. Cullis. 2021b. Protective Effect of Edaravone against Cationic Lipid-Mediated Oxidative Stress and Apoptosis. Biol Pharm Bull. 44:144–149.

157. Tilney LG, Bryan J, Bush DJ, Fujiwara K, Mooseker MS, Murphy DB, & DH Snyder (1973) Microtubules: evidence for 13 protofilaments. J Cell Biol. 59: 267–275.

158. Traub, P., C. Bauer, R. Hartig, S. Grub, and J. Stahl. 1998. Colocalization of single ribosomes with intermediate filaments in puromycin-treated and serum-starved mouse embryo fibroblasts. Biol Cell. 90:319–337.

159. Trylska, J., R. Konecny, F. Tama, C.L. Brooks, 3rd, and J.A. McCammon. 2004. Ribosome motions modulate electrostatic properties. Biopolymers. 74:423–431.

160. Unwin, P.N., and R.A. Milligan. 1982. A large particle associated with the perimeter of the nuclear pore complex. J Cell Biol. 93:63–75.

161. Varkouhi, A.K., M. Scholte, G. Storm, and H.J. Haisma. 2011. Endosomal escape pathways for delivery of biologicals. J Control Release. 151:220–228.

162. Venable, J.H., and R. Coggeshall. 1965. A Simplified Lead Citrate Stain for Use in Electron Microscopy. J Cell Biol. 25:407–408.

163. Verma, A., O. Uzun, Y. Hu, Y. Hu, H.S. Han, N. Watson, S. Chen, D.J. Irvine, and F. Stellacci. 2008. Surface-structure-regulated cell-membrane penetration by monolayer-protected nanoparticles. Nat Mater. 7:588–595.

164. Voigt, T., W. Dauber, I. Bensemann-Ryvkin, and X. Hartel. 2002. Increasing membrane contrast by means of imidazole-osmium post-fixation as exemplified by skeletal muscle fiber. Microsc Res Tech. 58:121–124.

165. Walter, R.J., and B. Tandler. 1989. Viruses and annulate lamellae in Friend erythroleukemia cells. J Submicrosc Cytol Pathol. 21:93–101.

166. Wan, C., T.M. Allen, and P.R. Cullis. 2014. Lipid nanoparticle delivery systems for siRNA-based therapeutics. Drug Deliv Transl Res. 4:74–83.

167. Wang, T., L.M. Larcher, L. Ma, and R.N. Veedu. 2018. Systematic Screening of Commonly Used Commercial Transfection Reagents towards Efficient Transfection of Single-Stranded Oligonucleotides. Molecules. 23.

168. Wheeler, R.J., and A.A. Hyman. 2018. Controlling compartmentalization by non-membrane-bound organelles. Philos Trans R Soc Lond B Biol Sci. 373.

169. Woodruff, J.B., A.A. Hyman, and E. Boke. 2018. Organization and Function of Non-dynamic Biomolecular Condensates. Trends Biochem Sci. 43:81–94.

170. Xu, Y., S.W. Hui, P. Frederik, and F.C. Szoka, Jr. 1999. Physicochemical characterization and purification of cationic lipoplexes. Biophys J. 77:341–353.

171. Yla-Anttila, P., H. Vihinen, E. Jokitalo, and E.L. Eskelinen. 2009. 3D tomography reveals connections between the phagophore and endoplasmic reticulum. Autophagy. 5:1180–1185.

172. Zabner, J., A.J. Fasbender, T. Moninger, K.A. Poellinger, and M.J. Welsh. 1995. Cellular and molecular barriers to gene transfer by a cationic lipid. J Biol Chem. 270:18997–19007.

173. Zhou, X., and L. Huang. 1994. DNA transfection mediated by cationic liposomes containing lipopolylysine: characterization and mechanism of action. Biochim Biophys Acta. 1189:195–203.

174. Zwilling, E., and F. Reggiori. 2022. Membrane Contact Sites in Autophagy. Cells. 11.

## REFERENCES (formatted)

175. Afzelius, B.A. 1992. Section staining for electron microscopy using tannic acid as a mordant: a simple method for visualization of glycogen and collagen. Microsc Res Tech. 21:65–72.

176. Alberti, S., and A.A. Hyman. 2021. Biomolecular condensates at the nexus of cellular stress, protein aggregation disease and ageing. Nat Rev Mol Cell Biol. 22:196–213.

177. Asensio, C.S., D.W. Sirkis, J.W. Maas, Jr., K. Egami, T.L. To, F.M. Brodsky, X. Shu, Y. Cheng, and R.H. Edwards. 2013. Self-assembly of VPS41 promotes sorting required for biogenesis of the regulated secretory pathway. Dev Cell. 27:425–437.

178. Aytar, B.S., J.P. Muller, S. Golan, S. Hata, H. Takahashi, Y. Kondo, Y. Talmon, N.L. Abbott, and D.M. Lynn. 2012. Addition of ascorbic acid to the extracellular environment activates lipoplexes of a ferrocenyl lipid and promotes cell transfection. J Control Release. 157:249–259.

179. Baker, N.A., D. Sept, S. Joseph, M.J. Holst, and J.A. McCammon. 2001. Electrostatics of nanosystems: application to microtubules and the ribosome. Proc Natl Acad Sci U S A. 98:10037–10041.

180. Ballarin-Gonzalez, B., and K.A. Howard. 2012. Polycation-based nanoparticle delivery of RNAi therapeutics: adverse effects and solutions. Adv Drug Deliv Rev. 64:1717–1729.

181. Banani, S.F., H.O. Lee, A.A. Hyman, and M.K. Rosen. 2017. Biomolecular condensates: organizers of cellular biochemistry. Nat Rev Mol Cell Biol. 18:285–298.

182. Basha, G., T.I. Novobrantseva, N. Rosin, Y.Y. Tam, I.M. Hafez, M.K. Wong, T. Sugo, V.M. Ruda, J. Qin, B. Klebanov, M. Ciufolini, A. Akinc, Y.K. Tam, M.J. Hope, and P.R. Cullis. 2011. Influence of cationic lipid composition on gene silencing properties of lipid nanoparticle formulations of siRNA in antigen-presenting cells. Mol Ther. 19:2186–2200.

183. Battersby, B.J., R. Grimm, S. Huebner, and G. Cevc. 1998. Evidence for three-dimensional interlayer correlations in cationic lipid-DNA complexes as observed by cryo-electron microscopy. Biochim Biophys Acta. 1372:379–383.

184. Bauer, C., and P. Traub. 1995. Interaction of intermediate filaments with ribosomes in vitro. Eur J Cell Biol. 68:288–296.

185. Bauer, M., B.W. Kristensen, M. Meyer, T. Gasser, H.R. Widmer, J. Zimmer, and M. Ueffing. 2006. Toxic effects of lipid-mediated gene transfer in ventral mesencephalic explant cultures. Basic Clin Pharmacol Toxicol. 98:395–400.

186. Bawdon, R.E., A.M. Fiskin, and R.G. Garrison. 1984. Ultrastructural aspects of cytoplasmic ribosomes from Histoplasma capsulatum and Blastomyces dermatitidis as revealed by heavy metal staining. Mycopathologia. 86:155–163.

187. Beiras, A., T. Garcia-Caballero, R. Gallego, and E. Roson. 1987. Staining of neuroendocrine Merkel cells of human epidermis using the uranaffin reaction. J Invest Dermatol. 89:366–368.

188. Bernhard, W. 1969. A new staining procedure for electron microscopical cytology. J Ultrastruct Res. 27:250–265.

189. Biazik, J., H. Vihinen, E. Jokitalo, and E.L. Eskelinen. 2017. Ultrastructural Characterization of Phagophores Using Electron Tomography on Cryoimmobilized and Freeze Substituted Samples. Methods Enzymol. 587:331–349.

190. Biazik, J., P. Yla-Anttila, H. Vihinen, E. Jokitalo, and E.L. Eskelinen. 2015. Ultrastructural relationship of the phagophore with surrounding organelles. Autophagy. 11:439–451.

191. Blad, D., L. Winqvist, and G. Dallner. 1977. Electrophoretic mobility of microsomes from rat liver. J Cell Sci. 23:285–297.

192. Blad-Holmberg, D. 1979. Study of electrophoretic mobility of cellular membranes isolated from rat liver. Biochim Biophys Acta. 553:25–39.

193. Boassa, D., M.L. Berlanga, M.A. Yang, M. Terada, J. Hu, E.A. Bushong, M. Hwang, E. Masliah, J.M. George, and M.H. Ellisman. 2013. Mapping the subcellular distribution of alpha-synuclein in neurons using genetically encoded probes for correlated light and electron microscopy: implications for Parkinson’s disease pathogenesis. J Neurosci. 33:2605–2615.

194. Borgese, N., N. Iacomino, S.F. Colombo, and F. Navone. 2021. The Link between VAPB Loss of Function and Amyotrophic Lateral Sclerosis. Cells. 10.

195. Brangwynne, C.P., C.R. Eckmann, D.S. Courson, A. Rybarska, C. Hoege, J. Gharakhani, F. Julicher, and A.A. Hyman. 2009. Germline P granules are liquid droplets that localize by controlled dissolution/condensation. Science. 324:1729–1732.

196. Brodie, D.A., P. Huie, M. Locke, and F.P. Ottensmeyer. 1982. The correlation between bismuth and uranyl staining and phosphorus content of intracellular structures as determined by electron spectroscopic imaging. Tissue Cell. 14:621–627.

197. Brunner, S., T. Sauer, S. Carotta, M. Cotten, M. Saltik, and E. Wagner. 2000. Cell cycle dependence of gene transfer by lipoplex, polyplex and recombinant adenovirus. Gene Ther. 7:401–407.

198. Caracciolo, G., R. Caminiti, M.A. Digman, E. Gratton, and S. Sanchez. 2009. Efficient escape from endosomes determines the superior efficiency of multicomponent lipoplexes. J Phys Chem B. 113:4995–4997.

199. Chernousova, S., and M. Epple. 2017. Live-cell imaging to compare the transfection and gene silencing efficiency of calcium phosphate nanoparticles and a liposomal transfection agent. Gene Ther. 24:282–289.

200. Colin, M., M. Maurice, G. Trugnan, M. Kornprobst, R.P. Harbottle, A. Knight, R.G. Cooper, A.D. Miller, J. Capeau, C. Coutelle, and M.C. Brahimi-Horn. 2000. Cell delivery, intracellular trafficking and expression of an integrin-mediated gene transfer vector in tracheal epithelial cells. Gene Ther. 7:139–152.

201. Cordes, V.C., S. Reidenbach, and W.W. Franke. 1996. Cytoplasmic annulate lamellae in cultured cells: composition, distribution, and mitotic behavior. Cell Tissue Res. 284:177–191.

202. Dalby, B., S. Cates, A. Harris, E.C. Ohki, M.L. Tilkins, P.J. Price, and V.C. Ciccarone. 2004. Advanced transfection with Lipofectamine 2000 reagent: primary neurons, siRNA, and high-throughput applications. Methods. 33:95–103.

203. Dallner, G., P. Siekevitz, and G.E. Palade. 1966. Biogenesis of endoplasmic reticulum membranes. I. Structural and chemical differentiation in developing rat hepatocyte. J Cell Biol. 30:73–96.

204. Damodaran, A.P., T. Courtheoux, E. Watrin, and C. Prigent. 2020. Alteration of SC35 localization by transfection reagents. Biochim Biophys Acta Mol Cell Res. 1867:118650.

205. Dokka, S., D. Toledo, X. Shi, V. Castranova, and Y. Rojanasakul. 2000. Oxygen radical-mediated pulmonary toxicity induced by some cationic liposomes. Pharm Res. 17:521–525.

206. Donkuru, M., I. Badea, S. Wettig, R. Verrall, M. Elsabahy, and M. Foldvari. 2010. Advancing nonviral gene delivery: lipid- and surfactant-based nanoparticle design strategies. Nanomedicine (Lond). 5:1103–1127.

207. Dunlap, D.D., A. Maggi, M.R. Soria, and L. Monaco. 1997. Nanoscopic structure of DNA condensed for gene delivery. Nucleic Acids Res. 25:3095–3101.

208. Duzgunes, N., and P.L. Felgner. 1993. Intracellular delivery of nucleic acids and transcription factors by cationic liposomes. Methods Enzymol. 221:303–306.

209. Duzgunes, N., J.A. Goldstein, D.S. Friend, and P.L. Felgner. 1989. Fusion of liposomes containing a novel cationic lipid, N-[2,3-(dioleyloxy)propyl]-N,N,N-trimethylammonium: induction by multivalent anions and asymmetric fusion with acidic phospholipid vesicles. Biochemistry. 28:9179–9184.

210. Dvorak, A.M., and E.S. Morgan. 2001. Ribosomes and secretory granules in human mast cells: close associations demonstrated by staining with a chelating agent. Immunol Rev. 179:94–101.

211. Erenpreisa, J., A. Ivanov, M. Cragg, G. Selivanova, and T. Illidge. 2002. Nuclear envelope-limited chromatin sheets are part of mitotic death. Histochem Cell Biol. 117:243–255.

212. Ewert, K., H.M. Evans, A. Ahmad, N.L. Slack, A.J. Lin, A. Martin-Herranz, and C.R. Safinya. 2005. Lipoplex structures and their distinct cellular pathways. Adv Genet. 53:119–155.

213. Ewert, K.K., P. Scodeller, L. Simon-Gracia, V.M. Steffes, E.A. Wonder, T. Teesalu, and C.R. Safinya. 2021. Cationic Liposomes as Vectors for Nucleic Acid and Hydrophobic Drug. Therapeutics. Pharmaceutics. 13.

214. Eymieux, S., Y. Rouille, O. Terrier, K. Seron, E. Blanchard, M. Rosa-Calatrava, J. Dubuisson, S. Belouzard, and P. Roingeard. 2021. Ultrastructural modifications induced by SARS-CoV-2 in Vero cells: a kinetic analysis of viral factory formation, viral particle morphogenesis and virion release. Cell Mol Life Sci. 78:3565–3576.

215. Farquhar, M.G., and G.E. Palade. 1965. Cell junctions in amphibian skin. J Cell Biol. 26:263–291.

216. Fasana, E., M. Fossati, A. Ruggiano, S. Brambillasca, C.C. Hoogenraad, F. Navone, M. Francolini, and N. Borgese. 2010. A VAPB mutant linked to amyotrophic lateral sclerosis generates a novel form of organized smooth endoplasmic reticulum. FASEB J. 24:1419–1430.

217. Felgner, J.H., R. Kumar, C.N. Sridhar, C.J. Wheeler, Y.J. Tsai, R. Border, P. Ramsey, M. Martin, and P.L. Felgner. 1994. Enhanced gene delivery and mechanism studies with a novel series of cationic lipid formulations. J Biol Chem. 269:2550–2561.

218. Felgner, P.L. 1997. Nonviral strategies for gene therapy. Sci Am. 276:102–106.

219. Felgner, P.L. 1999. Prospects for synthetic self-assembling systems in gene delivery. J Gene Med. 1:290–292.

220. Felgner, P.L., T.R. Gadek, M. Holm, R. Roman, H.W. Chan, M. Wenz, J.P. Northrop, G.M. Ringold, and M. Danielsen. 1987. Lipofection: a highly efficient, lipid-mediated DNA-transfection procedure. Proc Natl Acad Sci U S A. 84:7413–7417.

221. Fengsrud, M., N. Roos, T. Berg, W. Liou, J.W. Slot, and P.O. Seglen. 1995. Ultrastructural and immunocytochemical characterization of autophagic vacuoles in isolated hepatocytes: effects of vinblastine and asparagine on vacuole distributions. Exp Cell Res. 221:504–519.

222. Fenske, D.B., and P.R. Cullis. 2005. Entrapment of small molecules and nucleic acid-based drugs in liposomes. Methods Enzymol. 391:7–40.

223. Filion, M.C., and N.C. Phillips. 1997. Toxicity and immunomodulatory activity of liposomal vectors formulated with cationic lipids toward immune effector cells. Biochim Biophys Acta. 1329:345–356.

224. Friend, D.S., D. Papahadjopoulos, and R.J. Debs. 1996. Endocytosis and intracellular processing accompanying transfection mediated by cationic liposomes. Biochim Biophys Acta. 1278:41–50.

225. Frost, B., R.L. Jacks, and M.I. Diamond. 2009. Propagation of tau misfolding from the outside to the inside of a cell. J Biol Chem. 284:12845–12852.

226. Goryaynov, A., and W. Yang. 2014. Role of molecular charge in nucleocytoplasmic transport. PLoS One. 9:e88792.

227. Gudmundsson, S.R., K.A. Kallio, H. Vihinen, E. Jokitalo, N. Ktistakis, and E.L. Eskelinen. 2022. Morphology of Phagophore Precursors by Correlative Light-Electron Microscopy. Cells. 11.

228. Gustafsson, J., G. Arvidson, G. Karlsson, and M. Almgren. 1995. Complexes between cationic liposomes and DNA visualized by cryo-TEM. Biochim Biophys Acta. 1235:305–312.

229. Haggag, A., and J. Gilloteaux. 2005. Unusual circular annulate lamellae in hepatocytes of Torpedo marmorata. Histol Histopathol. 20:785–789.

230. Hanson, H.H., S. Kang, M. Fernandez-Monreal, T. Oung, M. Yildirim, R. Lee, K. Suyama, R.B. Hazan, and G.R. Phillips. 2010. LC3-dependent intracellular membrane tubules induced by gamma-protocadherins A3 and B2: a role for intraluminal interactions. J Biol Chem. 285:20982–20992.

231. Hawley-Nelson, P., and V. Ciccarone. 2001. Transfection of cultured eukaryotic cells using cationic lipid reagents. Curr Protoc Neurosci. Appendix 1:Appendix 1F.

232. Hawley-Nelson, P., and V. Ciccarone. 2003. Transfection of cultured eukaryotic cells using cationic lipid reagents. Curr Protoc Cell Biol. Chapter 20:Unit 20 26.

233. Hawley-Nelson, P., V. Ciccarone, and M.L. Moore. 2008. Transfection of cultured eukaryotic cells using cationic lipid reagents. Curr Protoc Mol Biol. Chapter 9:Unit 9 4.

234. Hayashi-Nishino, M., N. Fujita, T. Noda, A. Yamaguchi, T. Yoshimori, and A. Yamamoto. 2009. A subdomain of the endoplasmic reticulum forms a cradle for autophagosome formation. Nat Cell Biol. 11:1433–1437.

235. Heuser, J. 1981. Preparing biological samples for stereomicroscopy by the quick-freeze, deep-etch, rotary-replication technique. Methods Cell Biol. 22:97–122.

236. Heuser, J. 1989a. Protocol for 3-D visualization of molecules on mica via the quick-freeze, deep-etch technique. J Electron Microsc Tech. 13:244–263.

237. Heuser, J. 2000a. The production of ’cell cortices’ for light and electron microscopy. Traffic. 1:545–552.

238. Heuser, J.E. 1983. Procedure for freeze-drying molecules adsorbed to mica flakes. J Mol Biol. 169:155–195.

239. Heuser, J.E. 1989b. Development of the quick-freeze, deep-etch, rotary-replication technique of sample preparation for 3-D electron microscopy. Prog Clin Biol Res. 295:71–83.

240. Heuser, J.E. 2000b. Membrane traffic in anaglyph stereo. Traffic. 1:35–37.

241. Heuser, J.E. 2011. The origins and evolution of freeze-etch electron microscopy. J Electron Microsc (Tokyo). 60 Suppl 1:S3–29.

242. Heuser, J.E. 2022. The Structural Basis of Long-Term Potentiation in Hippocampal Synapses, Revealed by Electron Microscopy Imaging of Lanthanum-Induced Synaptic Vesicle Recycling. Front Cell Neurosci. 16:920360.

243. Heuser, J.E., T.S. Reese, M.J. Dennis, Y. Jan, L. Jan, and L. Evans. 1979. Synaptic vesicle exocytosis captured by quick freezing and correlated with quantal transmitter release. J Cell Biol. 81:275–300.

244. Heuser, J.E., and S.R. Salpeter. 1979. Organization of acetylcholine receptors in quick-frozen, deep-etched, and rotary-replicated Torpedo postsynaptic membrane. J Cell Biol. 82:150–173.

245. Heuser, J.E., and T.I. Tenkova. 2020. Introducing a mammalian nerve-muscle preparation ideal for physiology and microscopy, the transverse auricular muscle in the ear of the mouse. Neuroscience. 439:80–105.

246. Hoekstra, D., J. Rejman, L. Wasungu, F. Shi, and I. Zuhorn. 2007. Gene delivery by cationic lipids: in and out of an endosome. Biochem Soc Trans. 35:68–71.

247. Holmes, B.B., S.L. DeVos, N. Kfoury, M. Li, R. Jacks, K. Yanamandra, M.O. Ouidja, F.M. Brodsky, J. Marasa, D.P. Bagchi, P.T. Kotzbauer, T.M. Miller, D. Papy-Garcia, and M.I. Diamond. 2013. Heparan sulfate proteoglycans mediate internalization and propagation of specific proteopathic seeds. Proc Natl Acad Sci U S A. 110:E3138–3147.

248. Huber, S., A. Bar, S. Epp, J. Schmuckli-Maurer, N. Eberhard, B.M. Humbel, A. Hemphill, and K. Woods. 2020. Recruitment of Host Nuclear Pore Components to the Vicinity of Theileria Schizonts. mSphere. 5.

249. Huebner, S., B.J. Battersby, R. Grimm, and G. Cevc. 1999. Lipid-DNA complex formation: reorganization and rupture of lipid vesicles in the presence of DNA as observed by cryoelectron microscopy. Biophys J. 76:3158–3166.

250. Huxley, H.E., and G. Zubay. 1961. Preferential staining of nucleic acid-containing structures for electron microscopy. J Biophys Biochem Cytol. 11:273–296.

251. Hyman, A.A., and C.P. Brangwynne. 2011. Beyond stereospecificity: liquids and mesoscale organization of cytoplasm. Dev Cell. 21:14–16.

252. Hyman, A.A., C.A. Weber, and F. Julicher. 2014. Liquid-liquid phase separation in biology. Annu Rev Cell Dev Biol. 30:39–58.

253. Imreh, G., and E. Hallberg. 2000. An integral membrane protein from the nuclear pore complex is also present in the annulate lamellae: implications for annulate lamella formation. Exp Cell Res. 259:180–190.

254. Kanekura, K., I. Nishimoto, S. Aiso, and M. Matsuoka. 2006. Characterization of amyotrophic lateral sclerosis-linked P56S mutation of vesicle-associated membrane protein-associated protein B (VAPB/ALS8). J Biol Chem. 281:30223–30233.

255. Kanekura, K., H. Suzuki, S. Aiso, and M. Matsuoka. 2009. ER stress and unfolded protein response in amyotrophic lateral sclerosis. Mol Neurobiol. 39:81–89.

256. Karnovsky, M.J. 1961. Simple methods for "staining with lead" at high pH in electron microscopy. J Biophys Biochem Cytol. 11:729–732.

257. Karnovsky, M.J. 1971. Use of ferrocyanide-reduced osmium tetroxide in electron microscopy . In Abstracts of the American Society of Cell Biology. Abstracts of the American Society of Cell Biology, New Orleans. 146.

258. Kawabuchi, M., M. Osame, Y. Aika, and T. Kanaseki. 1989. Annulate lamellae-soleplate nuclei associations in skeletal muscle fibers of rats during chronic high-dose exposure to neostigmine. Anat Rec. 225:1–10.

259. Kessel, R.G. 1983a. Fibrogranular bodies, annulate lamellae, and polyribosomes in the dragonfly oocyte. J Morphol. 176:171–180.

260. Kessel, R.G. 1983b. The structure and function of annulate lamellae: porous cytoplasmic and intranuclear membranes. Int Rev Cytol. 82:181–303.

261. Kfoury, N., B.B. Holmes, H. Jiang, D.M. Holtzman, and M.I. Diamond. 2012. Trans-cellular propagation of Tau aggregation by fibrillar species. J Biol Chem. 287:19440–19451.

262. Kislev, N., H. Swift, and L. Bogorad. 1965. Nucleic Acids of Chloroplasts and Mitochondria in Swiss Chard. J Cell Biol. 25:327–344.

263. Knight, A.M., P.H. Culviner, N. Kurt-Yilmaz, T. Zou, S.B. Ozkan, and S. Cavagnero. 2013. Electrostatic effect of the ribosomal surface on nascent polypeptide dynamics. ACS Chem Biol. 8:1195–1204.

264. Kolay, S., and M.I. Diamond. 2020. Alzheimer’s disease risk modifier genes do not affect tau aggregate uptake, seeding or maintenance in cell models. FEBS Open Bio. 10:1912–1920.

265. Koltover, I., T. Salditt, J.O. Radler, and C.R. Safinya. 1998. An inverted hexagonal phase of cationic liposome-DNA complexes related to DNA release and delivery. Science. 281:78–81.

266. Konopka, K., G.S. Harrison, P.L. Felgner, and N. Duzgunes. 1997. Cationic liposome-mediated expression of HIV-regulated luciferase and diphtheria toxin a genes in HeLa cells infected with or expressing HIV. Biochim Biophys Acta. 1356:185–197.

267. Konopka, K., E. Pretzer, P.L. Felgner, and N. Duzgunes. 1996. Human immunodeficiency virus type-1 (HIV-1) infection increases the sensitivity of macrophages and THP-1 cells to cytotoxicity by cationic liposomes. Biochim Biophys Acta. 1312:186–196.

268. Korn, A.P., P. Spitnik-Elson, and D. Elson. 1985. Identification of the collar-like structure of the 30S ribosomal subunit from E. coli by dark field electron microscopy. Eur J Cell Biol. 39:56–61.

269. Kovacs, A.L., A. Reith, and P.O. Seglen. 1982. Accumulation of autophagosomes after inhibition of hepatocytic protein degradation by vinblastine, leupeptin or a lysosomotropic amine. Exp Cell Res. 137:191–201.

270. Kuijpers, M., V. van Dis, E.D. Haasdijk, M. Harterink, K. Vocking, J.A. Post, W. Scheper, C.C. Hoogenraad, and D. Jaarsma. 2013. Amyotrophic lateral sclerosis (ALS)-associated VAPB-P56S inclusions represent an ER quality control compartment. Acta Neuropathol Commun. 1:24.

271. Kulkarni, J.A., M.M. Darjuan, J.E. Mercer, S. Chen, R. van der Meel, J.L. Thewalt, Y.Y.C. Tam, and P.R. Cullis. 2018. On the Formation and Morphology of Lipid Nanoparticles Containing Ionizable Cationic Lipids and siRNA. ACS Nano. 12:4787–4795.

272. Kulkarni, J.A., S.B. Thomson, J. Zaifman, J. Leung, P.K. Wagner, A. Hill, Y.Y.C. Tam, P.R. Cullis, T.L. Petkau, and B.R. Leavitt. 2020. Spontaneous, solvent-free entrapment of siRNA within lipid nanoparticles. Nanoscale. 12:23959–23966.

273. Kulkarni, J.A., D. Witzigmann, S. Chen, P.R. Cullis, and R. van der Meel. 2019a. Lipid Nanoparticle Technology for Clinical Translation of siRNA Therapeutics. Acc Chem Res. 52:2435–2444.

274. Kulkarni, J.A., D. Witzigmann, J. Leung, R. van der Meel, J. Zaifman, M.M. Darjuan, H.M. Grisch-Chan, B. Thony, Y.Y.C. Tam, and P.R. Cullis. 2019b. Fusion-dependent formation of lipid nanoparticles containing macromolecular payloads. Nanoscale. 11:9023–9031.

275. Kuntsche, J., J.C. Horst, and H. Bunjes. 2011. Cryogenic transmission electron microscopy (cryo-TEM) for studying the morphology of colloidal drug delivery systems. Int J Pharm. 417:120–137.

276. Lazebnik, M., R.K. Keswani, and D.W. Pack. 2016. Endocytic Transport of Polyplex and Lipoplex siRNA Vectors in HeLa Cells. Pharm Res. 33:2999–3011.

277. Le Bihan, O., R. Chevre, S. Mornet, B. Garnier, B. Pitard, and O. Lambert. 2011. Probing the in vitro mechanism of action of cationic lipid/DNA lipoplexes at a nanometric scale. Nucleic Acids Res. 39:1595–1609.

278. Leal, C., N.F. Bouxsein, K.K. Ewert, and C.R. Safinya. 2010. Highly efficient gene silencing activity of siRNA embedded in a nanostructured gyroid cubic lipid matrix. J Am Chem Soc. 132:16841–16847.

279. Leung, A.K., Y.Y. Tam, S. Chen, I.M. Hafez, and P.R. Cullis. 2015. Microfluidic Mixing: A General Method for Encapsulating Macromolecules in Lipid Nanoparticle Systems. J Phys Chem B. 119:8698–8706.

280. Li, Z., C. Zhang, Z. Wang, J. Shen, P. Xiang, X. Chen, J. Nan, and Y. Lin. 2019. Lipofectamine 2000/siRNA complexes cause endoplasmic reticulum unfolded protein response in human endothelial cells. J Cell Physiol. 234:21166–21181.

281. Lin, A.J., N.L. Slack, A. Ahmad, C.X. George, C.E. Samuel, and C.R. Safinya. 2003. Three-dimensional imaging of lipid gene-carriers: membrane charge density controls universal transfection behavior in lamellar cationic liposome-DNA complexes. Biophys J. 84:3307–3316.

282. Lin, D.H., and A. Hoelz. 2019. The Structure of the Nuclear Pore Complex (An Update). Annu Rev Biochem. 88:725–783.

283. Lin, Q., J. Chen, Z. Zhang, and G. Zheng. 2014. Lipid-based nanoparticles in the systemic delivery of siRNA. Nanomedicine (Lond). 9:105–120.

284. Liu, Y., L. Wenning, M. Lynch, and T.M. Reineke. 2004. New poly(d-glucaramidoamine)s induce DNA nanoparticle formation and efficient gene delivery into mammalian cells. J Am Chem Soc. 126:7422–7423.

285. Lozhkin, A., A.E. Vendrov, H. Pan, S.A. Wickline, N.R. Madamanchi, and M.S. Runge. 2017. NADPH oxidase 4 regulates vascular inflammation in aging and atherosclerosis. J Mol Cell Cardiol. 102:10–21.

286. Luk, K.C., C. Song, P. O’Brien, A. Stieber, J.R. Branch, K.R. Brunden, J.Q. Trojanowski, and V.M. Lee. 2009. Exogenous alpha-synuclein fibrils seed the formation of Lewy body-like intracellular inclusions in cultured cells. Proc Natl Acad Sci U S A. 106:20051–20056.

287. Lv, H., S. Zhang, B. Wang, S. Cui, and J. Yan. 2006. Toxicity of cationic lipids and cationic polymers in gene delivery. J Control Release. 114:100–109.

288. Magalhaes, S., S. Duarte, G.A. Monteiro, and F. Fernandes. 2014. Quantitative evaluation of DNA dissociation from liposome carriers and DNA escape from endosomes during lipid-mediated gene delivery. Hum Gene Ther Methods. 25:303–313.

289. Mahamid, J., S. Pfeffer, M. Schaffer, E. Villa, R. Danev, L.K. Cuellar, F. Forster, A.A. Hyman, J.M. Plitzko, and W. Baumeister. 2016. Visualizing the molecular sociology at the HeLa cell nuclear periphery. Science. 351:969–972.

290. Majzoub, R.N., C.L. Chan, K.K. Ewert, B.F. Silva, K.S. Liang, and C.R. Safinya. 2015. Fluorescence microscopy colocalization of lipid-nucleic acid nanoparticles with wildtype and mutant Rab5-GFP: A platform for investigating early endosomal events. Biochim Biophys Acta. 1848:1308–1318.

291. Majzoub, R.N., K.K. Ewert, and C.R. Safinya. 2016a. Cationic liposome-nucleic acid nanoparticle assemblies with applications in gene delivery and gene silencing. Philos Trans A Math Phys Eng Sci. 374.

292. Majzoub, R.N., K.K. Ewert, and C.R. Safinya. 2016b. Quantitative Intracellular Localization of Cationic Lipid-Nucleic Acid Nanoparticles with Fluorescence Microscopy. Methods Mol Biol. 1445:77–108.

293. Malatesta, M. 2021. Transmission Electron Microscopy as a Powerful Tool to Investigate the Interaction of Nanoparticles with Subcellular Structures. Int J Mol Sci. 22.

294. Malone, R.W., P.L. Felgner, and I.M. Verma. 1989. Cationic liposome-mediated RNA transfection. Proc Natl Acad Sci U S A. 86:6077–6081.

295. Martin, T.M., B.J. Wysocki, J.P. Beyersdorf, T.A. Wysocki, and A.K. Pannier. 2014. Integrating mitosis, toxicity, and transgene expression in a telecommunications packet-switched network model of lipoplex-mediated gene delivery. Biotechnol Bioeng. 111:1659–1671.

296. Maupin P, & TD Pollard (1983) Improved preservation and staining of cytoplasmic structures by tannic acid-glutaraldehyde-saponin fixation. J Cell Biol. 96: 51–62.

297. Maurer, N., A. Mori, L. Palmer, M.A. Monck, K.W. Mok, B. Mui, Q.F. Akhong, and P.R. Cullis. 1999. Lipid-based systems for the intracellular delivery of genetic drugs. Mol Membr Biol. 16:129–140.

298. Midoux, P., C. Pichon, J.J. Yaouanc, and P.A. Jaffres. 2009. Chemical vectors for gene delivery: a current review on polymers, peptides and lipids containing histidine or imidazole as nucleic acids carriers. Br J Pharmacol. 157:166–178.

299. Mo, R.H., J.L. Zaro, J.H. Ou, and W.C. Shen. 2012. Effects of Lipofectamine 2000/siRNA complexes on autophagy in hepatoma cells. Mol Biotechnol. 51:1–8.

300. Morgenstern, E. 1977. [Cytochemical studies of ultrathin frozen sections of blood platelets]. Acta Histochem Suppl. Suppl 18:259–264.

301. Napoli, E., S. Liu, I. Marsilio, K. Zarbalis, and C. Giulivi. 2017. Lipid-based DNA/siRNA transfection agents disrupt neuronal bioenergetics and mitophagy. Biochem J. 474:3887–3902.

302. Navone, F., P. Genevini, and N. Borgese. 2015. Autophagy and Neurodegeneration: Insights from a Cultured Cell Model of ALS. Cells. 4:354–386.

303. Nekooki-Machida, Y., M. Kurosawa, N. Nukina, K. Ito, T. Oda, and M. Tanaka. 2009. Distinct conformations of in vitro and in vivo amyloids of huntingtin-exon1 show different cytotoxicity. Proc Natl Acad Sci U S A. 106:9679–9684.

304. Nishimura, A.L., M. Mitne-Neto, H.C. Silva, A. Richieri-Costa, S. Middleton, D. Cascio, F. Kok, J.R. Oliveira, T. Gillingwater, J. Webb, P. Skehel, and M. Zatz. 2004. A mutation in the vesicle-trafficking protein VAPB causes late-onset spinal muscular atrophy and amyotrophic lateral sclerosis. Am J Hum Genet. 75:822–831.

305. Niso-Santano, M., J.M. Bravo-San Pedro, R. Gomez-Sanchez, V. Climent, G. Soler, J.M. Fuentes, and R.A. Gonzalez-Polo. 2011. ASK1 overexpression accelerates paraquat-induced autophagy via endoplasmic reticulum stress. Toxicol Sci. 119:156–168.

306. Ohki, E.C., M.L. Tilkins, V.C. Ciccarone, and P.J. Price. 2001. Improving the transfection efficiency of post-mitotic neurons. J Neurosci Methods. 112:95–99.

307. Papiani, G., A. Ruggiano, M. Fossati, A. Raimondi, G. Bertoni, M. Francolini, R. Benfante, F. Navone, and N. Borgese. 2012. Restructured endoplasmic reticulum generated by mutant amyotrophic lateral sclerosis-linked VAPB is cleared by the proteasome. J Cell Sci. 125:3601–3611.

308. Park, S., C. Zuber, and J. Roth. 2020. Selective autophagy of cytosolic protein aggregates involves ribosome-free rough endoplasmic reticulum. Histochem Cell Biol. 153:89–99.

309. Payne, C.M., R.B. Nagle, V.F. Borduin, and A. Kim. 1983. An ultrastructural evaluation of the cell organelle specificity of the uranaffin reaction in two human endocrine neoplasms. J Submicrosc Cytol. 15:833–841.

310. Piefke, J., T. Arad, H.S. Gewitz, A. Yonath, and H.G. Wittmann. 1986. The growth of ordered two-dimensional sheets of 70 S ribosomes from Bacillus stearothermophilus. FEBS Lett. 209:104–106.

311. Plaza-Ga, I., V. Manzaneda-Gonzalez, M. Kisovec, V. Almendro-Vedia, M. Munoz-Ubeda, G. Anderluh, A. Guerrero-Martinez, P. Natale, and I. Lopez Montero. 2019. pH-triggered endosomal escape of pore-forming Listeriolysin O toxin-coated gold nanoparticles. J Nanobiotechnology. 17:108.

312. Radler, J.O., I. Koltover, T. Salditt, and C.R. Safinya. 1997. Structure of DNA-cationic liposome complexes: DNA intercalation in multilamellar membranes in distinct interhelical packing regimes. Science. 275:810–814.

313. Raghunayakula, S., D. Subramonian, M. Dasso, R. Kumar, and X.D. Zhang. 2015. Molecular Characterization and Functional Analysis of Annulate Lamellae Pore Complexes in Nuclear Transport in Mammalian Cells. PLoS One. 10:e0144508.

314. Ramezanpour, M., M.L. Schmidt, I. Bodnariuc, J.A. Kulkarni, S.S.W. Leung, P.R. Cullis, J.L. Thewalt, and D.P. Tieleman. 2019. Ionizable amino lipid interactions with POPC: implications for lipid nanoparticle function. Nanoscale. 11:14141–14146.

315. Reifarth, M., S. Hoeppener, and U.S. Schubert. 2018. Uptake and Intracellular Fate of Engineered Nanoparticles in Mammalian Cells: Capabilities and Limitations of Transmission Electron Microscopy-Polymer-Based Nanoparticles. Adv Mater. 30.

316. Reynolds, E.S. 1963. The use of lead citrate at high pH as an electron-opaque stain in electron microscopy. J Cell Biol. 17:208–212.

317. Safinya, C.R., K.K. Ewert, R.N. Majzoub, and C. Leal. 2014. Cationic liposome-nucleic acid complexes for gene delivery and gene silencing. New J Chem. 38:5164–5172.

318. Sanders, D.W., S.K. Kaufman, S.L. DeVos, A.M. Sharma, H. Mirbaha, A. Li, S.J. Barker, A.C. Foley, J.R. Thorpe, L.C. Serpell, T.M. Miller, L.T. Grinberg, W.W. Seeley, and M.I. Diamond. 2014. Distinct tau prion strains propagate in cells and mice and define different tauopathies. Neuron. 82:1271–1288.

319. Sawaguchi, A., S. Ide, Y. Goto, J. Kawano, T. Oinuma, and T. Suganuma. 2001. A simple contrast enhancement by potassium permanganate oxidation for Lowicryl K4M ultrathin sections prepared by high pressure freezing/freeze substitution. J Microsc. 201:77–83.

320. Schavemaker, P.E., W.M. Smigiel, and B. Poolman. 2017. Ribosome surface properties may impose limits on the nature of the cytoplasmic proteome. Elife. 6.

321. Schiavone, N., L. Papucci, P. Luciani, A. Lapucci, M. Donnini, and S. Capaccioli. 2000. Induction of apoptosis and mitosis inhibition by degraded DNA lipotransfection mimicking genotoxic drug effects. Biochem Biophys Res Commun. 270:406–414.

322. Shu, X., V. Lev-Ram, T.J. Deerinck, Y. Qi, E.B. Ramko, M.W. Davidson, Y. Jin, M.H. Ellisman, and R.Y. Tsien. 2011. A genetically encoded tag for correlated light and electron microscopy of intact cells, tissues, and organisms. PLoS Biol. 9:e1001041.

323. Silva, J.P., I.M. Oliveira, A.C. Oliveira, M. Lucio, A.C. Gomes, P.J. Coutinho, and M.E. Oliveira. 2014. Structural dynamics and physicochemical properties of pDNA/DODAB:MO lipoplexes: effect of pH and anionic lipids in inverted non-lamellar phases versus lamellar phases. Biochim Biophys Acta. 1838:2555–2567.

324. Singh, R.S., C. Goncalves, P. Sandrin, C. Pichon, P. Midoux, and A. Chaudhuri. 2004. On the gene delivery efficacies of pH-sensitive cationic lipids via endosomal protonation: a chemical biology investigation. Chem Biol. 11:713–723.

325. Soloff, B.L. 1973. Buffered potassium permanganate-uranyl acetate-lead citrate staining seguence for ultrathin sections. Stain Technol. 48:159–165.

326. Stephan, D.J., Z.Y. Yang, H. San, R.D. Simari, C.J. Wheeler, P.L. Felgner, D. Gordon, G.J. Nabel, and E.G. Nabel. 1996. A new cationic liposome DNA complex enhances the efficiency of arterial gene transfer in vivo. Hum Gene Ther. 7:1803–1812.

327. Sternberg, B., K. Hong, W. Zheng, and D. Papahadjopoulos. 1998. Ultrastructural characterization of cationic liposome-DNA complexes showing enhanced stability in serum and high transfection activity in vivo. Biochim Biophys Acta. 1375:23–35.

328. Sternberg, B., F.L. Sorgi, and L. Huang. 1994. New structures in complex formation between DNA and cationic liposomes visualized by freeze-fracture electron microscopy. FEBS Lett. 356:361–366.

329. Terada, T., J.A. Kulkarni, A. Huynh, S. Chen, R. van der Meel, Y.Y.C. Tam, and P.R. Cullis. 2021a. Characterization of Lipid Nanoparticles Containing Ionizable Cationic Lipids Using Design-of-Experiments Approach. Langmuir. 37:1120–1128.

330. Terada, T., J.A. Kulkarni, A. Huynh, Y.Y.C. Tam, and P. Cullis. 2021b. Protective Effect of Edaravone against Cationic Lipid-Mediated Oxidative Stress and Apoptosis. Biol Pharm Bull. 44:144–149.

331. Tilney LG, Bryan J, Bush DJ, Fujiwara K, Mooseker MS, Murphy DB, & DH Snyder (1973) Microtubules: evidence for 13 protofilaments. J Cell Biol. 59: 267–275.

332. Traub, P., C. Bauer, R. Hartig, S. Grub, and J. Stahl. 1998. Colocalization of single ribosomes with intermediate filaments in puromycin-treated and serum-starved mouse embryo fibroblasts. Biol Cell. 90:319–337.

333. Trylska, J., R. Konecny, F. Tama, C.L. Brooks, 3rd, and J.A. McCammon. 2004. Ribosome motions modulate electrostatic properties. Biopolymers. 74:423–431.

334. Unwin, P.N., and R.A. Milligan. 1982. A large particle associated with the perimeter of the nuclear pore complex. J Cell Biol. 93:63–75.

335. Varkouhi, A.K., M. Scholte, G. Storm, and H.J. Haisma. 2011. Endosomal escape pathways for delivery of biologicals. J Control Release. 151:220–228.

336. Venable, J.H., and R. Coggeshall. 1965. A Simplified Lead Citrate Stain for Use in Electron Microscopy. J Cell Biol. 25:407–408.

337. Verma, A., O. Uzun, Y. Hu, Y. Hu, H.S. Han, N. Watson, S. Chen, D.J. Irvine, and F. Stellacci. 2008. Surface-structure-regulated cell-membrane penetration by monolayer-protected nanoparticles. Nat Mater. 7:588–595.

338. Voigt, T., W. Dauber, I. Bensemann-Ryvkin, and X. Hartel. 2002. Increasing membrane contrast by means of imidazole-osmium post-fixation as exemplified by skeletal muscle fiber. Microsc Res Tech. 58:121–124.

339. Walter, R.J., and B. Tandler. 1989. Viruses and annulate lamellae in Friend erythroleukemia cells. J Submicrosc Cytol Pathol. 21:93–101.

340. Wan, C., T.M. Allen, and P.R. Cullis. 2014. Lipid nanoparticle delivery systems for siRNA-based therapeutics. Drug Deliv Transl Res. 4:74–83.

341. Wang, T., L.M. Larcher, L. Ma, and R.N. Veedu. 2018. Systematic Screening of Commonly Used Commercial Transfection Reagents towards Efficient Transfection of Single-Stranded Oligonucleotides. Molecules. 23.

342. Wheeler, R.J., and A.A. Hyman. 2018. Controlling compartmentalization by non-membrane-bound organelles. Philos Trans R Soc Lond B Biol Sci. 373.

343. Woodruff, J.B., A.A. Hyman, and E. Boke. 2018. Organization and Function of Non-dynamic Biomolecular Condensates. Trends Biochem Sci. 43:81–94.

344. Xu, Y., S.W. Hui, P. Frederik, and F.C. Szoka, Jr. 1999. Physicochemical characterization and purification of cationic lipoplexes. Biophys J. 77:341–353.

345. Yla-Anttila, P., H. Vihinen, E. Jokitalo, and E.L. Eskelinen. 2009. 3D tomography reveals connections between the phagophore and endoplasmic reticulum. Autophagy. 5:1180–1185.

346. Zabner, J., A.J. Fasbender, T. Moninger, K.A. Poellinger, and M.J. Welsh. 1995. Cellular and molecular barriers to gene transfer by a cationic lipid. J Biol Chem. 270:18997–19007.

347. Zhou, X., and L. Huang. 1994. DNA transfection mediated by cationic liposomes containing lipopolylysine: characterization and mechanism of action. Biochim Biophys Acta. 1189:195–203.

348. Zwilling, E., and F. Reggiori. 2022. Membrane Contact Sites in Autophagy. Cells. 11.

